# Appetitive cue exposure increases neural reward responses without modulating temporal discounting

**DOI:** 10.1101/2022.11.21.517327

**Authors:** Kilian Knauth, David Mathar, Bojana Kuzmanovic, Marc Tittgemeyer, Jan Peters

## Abstract

When given a choice, humans and many animals prefer smaller but sooner over larger but later rewards, a tendency referred to as temporal discounting. Alterations in devaluation of future rewards have been reported in a range of maladaptive behaviors and clinical conditions. Although temporal discounting is highly stable over time and testing environments (e.g., laboratory vs. virtual reality), it is partly under contextual control. For example, highly appetitive cues such as erotic images might increase preferences for immediate rewards, although overall evidence remains mixed. Dopaminergic circuit activity and striatal dopamine concentrations are often assumed to drive increases in temporal discounting following appetitive cue-exposure, yet this was never explicitly tested. Here we examined cue-reactivity effects (erotic vs. neutral pictures) on subsequent temporal discounting in a pre-registered within-subjects study in healthy male participants (n=38). Functional magnetic resonance imaging (fMRI) assessed neural cue-reactivity, value-computations and choice-related effects. Preregistered analyses replicated previous findings of value coding in ventromedial prefrontal cortices, striatum and cingulate cortex. Likewise, as hypothesized, lateral prefrontal cortex activity increased during choices of delayed rewards, potentially reflecting cognitive control. As predicted, erotic vs. neutral cue exposure was associated with increased activity in attention and reward circuits. Contrary to our preregistered hypotheses, temporal discounting was largely unaffected by cue exposure. Likewise, cue-reactivity in key areas of the dopaminergic reward circuit (Nacc, VTA) was not significantly associated with changes in behavior. Our results indicate that behavioral effects of erotic cue exposure on temporal discounting might not be as unequivocal as previously thought and raise doubt on the hypothesis of an upregulated dopaminergic ramping mechanism, that might support myopic approach behavior towards immediate rewards.

## Introduction

Intertemporal decisions require a trade-off between smaller, sooner (SS) and larger, later (LL) rewards. People and many animals tend to discount the value of the later rewards as a function of time, resulting in an increased preference for immediate rewards (temporal discounting (TD); Kalenscher & Pennertz, 2008; Peters & Büchel, 2011). The degree of TD varies substantially across individuals (Peters & Büchel, 2011; Soman et al., 2005). High discount rates are observed in clinical groups exhibiting impulsive and/or short-sighted behavior (Bulley & Schacter, 2020), including gambling disorder, substance abuse, impulse control disorders or lesions to the prefrontal cortices (Amlung et al., 2019; Garofalo et al., 2022; Lempert et al., 2019; Peters & D’Esposito, 2016; Weinsztok et al., 2021). TD is a largely stable trait, with long-term test-retest reliabilities ranging from .71 to .91 (Arfer & Luhmann, 2017; Bruder et al., 2021; Enkavi et al., 2019; Kirby et al., 2009; Simpson & Vuchinich, 2000).

However, TD is additionally affected by environmental factors and cues (Lempert & Phelps, 2016; Peters & Büchel, 2011). In men, TD increases following block-wise presentation of arousing images of opposite-sex faces or erotica (Kim & Zauberman, 2013; Van den Bergh et al., 2008; Wilson & Dali, 2004), stimuli which possess inherently rewarding or appetitive qualities and elicit basic emotional responses (Klucken et al., 2013). More recent results support a more fine-graded association between visual appetitive stimulus processing and impulsivity, possibly moderated by internal motivational or metabolic conditions (Chio et al., 2015; Otterbring & Sela, 2020). Chiou and colleagues (2015) found that a *mating mindset* mediated the connection between viewing pictures of attractive women and greater TD in men. Otterbring and Sela (2020) reported more impatient financial decisions following sexually arousing ads compared to neutral - but only in hungry male individuals. Such internal states might foster active approach behavior towards immediate rewards.

Previous studies hypothesized that an upregulation of reward circuitry following appetitive cue exposure might drive this effect (Van den Bergh et al., 2008). Indeed, exposure to primary reinforcers including appetitive (erotic) cues increases activity in reward circuits, including ventral striatum (VS), orbitofrontal cortex (OFC) and ventral tegmental area (VTA; Gola et al., 2016; Markert et al., 2021; Stark et al., 2019; Wehrum-Osinsky et al., 2014). Such exposure might also lead to a bias towards short-term rewards (Li, 2008; Mathar et al., 2022; Yeomans & Brace, 2015) possibly driven by increased dopamine release. Cortical and striatal dopamine tone can modulate TD (Arrondo et al., 2015; Cools, 2008; de Wit, 2002; Hamidovic et al., 2008; Kayser et al., 2012; Petzold et al., 2019; Pine et al., 2010; Wagner et al., 2020; Weber et al., 2016). While the directionality of these effects is mixed, overall evidence appears to be more compatible with a reduction in TD following increases in dopamine tone (D’Armour-Horvat & Leyton, 2014).

Erotic cue processing and a resulting present-orientation in healthy participants might share conceptual similarities with cue-reactivity in addiction, referring to increased subjective, physiological and neural responses to addiction-related cues (Courtney et al., 2015; Starcke et al., 2018; Volkow et al., 2010; Zhou et al., 2019). Exposure to addiction-related cues can drive increases in TD in gambling disorder (Dixon et al., 2006; Miedl et al., 2014; Wagner et al., 2022). The study of appetitive cue effects on TD in healthy participants might thus inform our understanding of maladaptive behaviors in clinical groups and potential interventions.

To sum up, there is considerable evidence that exposure to highly appetitive (erotic) cues can increase TD (Kim & Zauberman, 2013; Otterbring et al., 2020; Wilson & Daly, 2004) and that erotic cues upregulate activity in reward-related (dopaminergic) regions (Gola et al., 2016; Stark et al., 2005; Stark et al., 2019; Wehrum-Osinsky et al., 2014). However, the degree to which neuronal (erotic) cue-reactivity in these areas directly contributes to changes in TD remains unclear.

The current study addressed these issues in the following ways. First, extending previous work, we used fMRI to directly measure the effects of erotic cue exposure on reward circuit activity and subsequent temporal discounting. Second, we linked reward-system-reactivity to TD. Based on the previous literature we preregistered the following hypotheses (https://osf.io/w5puk/): On the behavioral level we expected increased TD following erotic cue exposure (H1). On the neuronal level we predicted to replicate previous findings on neural erotic cue processing, subjective value coding and choice-related activity. Specifically, we predicted subjective value coding of delayed rewards in striatum and ventromedial prefrontal cortex (vmPFC) (H2; Peters & Büchel, 2009) and increased lateral prefrontal cortex activity (lPFC) during choices of LL vs. SS rewards (H3; Smith et al., 2018). Further, we expected erotic vs. neutral cues to upregulate activity in a set of a priori-defined regions related to the processing of visual erotic stimuli (H4; Stark et al., 2019, see methods section for a detailed procedure on ROI definition). To link neuronal effects to TD, we tested two predictions. First, lPFC activity at LL-option onset was predicted to be reduced following erotic vs. neutral cues (H5). Second, we predicted increased reward-system-reactivity (erotic > neutral) within key dopaminergic regions (Nacc, VTA) and reduced LPFC activity in response to erotic cues to be positively associated with cue-induced increases in TD (H6).

## 2. Materials and Methods

### 2.1. Participants

Based on mean effect size estimates from two previous studies on erotic cue exposure effects on TD (Kim and Zauberman, 2013; Wilson & Daly, 2004), a power analysis (G*Power; Faul et al., 2007) yielded a preregistered sample size of N = 31 when taking a test-retest reliability estimate of TD into account (Enkavi et al., 2019) (effect size Cohen’s f = 0.22, error probability α = .05, power = .80; F-tests, number of groups: 1; number of measurements: 2; correlation between repeated measures: 0.65). To account for potential drop out and data loss, we tested a total sample of 38 participants. Two participants dropped out after the first testing session. fMRI data from one additional participant was lost due to technical error at the MRI environment, while behavioral data was preserved. The final sample therefore consisted of N = 36 male participants (mean age ± SD (range) = 31.2 ± 7.5 (20-50)). Participants were recruited via advertisements on internet bulletin boards, mailing lists and local notices. Main inclusion criteria included male gender, right-handedness, heterosexuality, normal or corrected-to-normal vision, no alcohol or drug abuse, no psychiatric, neurological or cardiovascular disease (past or current) and no pacemakers or other ferromagnetic materials on the body. All experimental procedures were approved by the institutional ethics committee of the University of Cologne Medical Center (application number: 17-045), and participants provided informed written consent prior to participation in the study.

### 2.2. Appetitive cues

During each of the fMRI sessions, participants underwent two analogous cue exposure phases and performed two different decision-making tasks (see Tasks & Procedure section 2.3). Depending on the experimental condition of the day, participants were either exposed to erotic or neutral visual stimuli. Experimental images were partly derived from IAPS database, Nencki Affective Picture System (NAPS), EmoPics (Lang, Bradley, & Cuthbert, 2008; Marchewka et al., 2014; Wessa et al., 2010) and from a google search. Our preliminary stimulus set consisted of 220 erotic and neutral images which were roughly matched for image content and complexity. In a preceding pilot study, the preliminary set was rated concerning valence and arousal levels by an independent sample. The most arousing erotic (N=90) and the least arousing neutral images (N=90) were included into our experimental image pool. Consequently, erotic and neutral cues differed in arousal (erotic: 65.07 ± 3.51, neutral: 4.89 ± 3.39; *t*_*(178)*_ = 140.67, *p* < 0.001) and valence (erotic: 64.92 ± 3.39, neutral: 48.90 ± 9.84; *t*_*(178)*_ = 14.59, *p* < 0.001). We ensured that images were matched on intensity (erotic: 0.46 ± 0.09, neutral: 0.45 ± 0.14; *t*_*(178)*_ = 0.26, *p* = 0.79) and contrast (erotic: 0.19 ± 0.04, neutral: 0.19 ± 0.03; *t*_*(178)*_ = -0.47, *p* = 0.64). Control scrambled images were created by randomly shuffling pixel locations, thereby preserving intensity and contrast. Unique image sets were created for each participant and for each cue phase by randomly drawing forty intact and twenty scrambled control images without replacement from their respective image pools (N=90).

### 2.3. Tasks & Procedure

The current study was conducted as one group within-subject design, including two experimental conditions (erotic vs. neutral). Data collection took place on two testing days with an approximate interval of 11 days (mean ± SD (range) = 11.31±12.62 (1-70)). Each day, participants performed two decision-making tasks and two cue-exposure phases during fMRI. After introduction to the experimental set-up and scanning-preparation, participants completed the first cue-exposure phase. The cue phase consisted of 40 neutral or appetitive (erotic) images (depending on the condition on that day) and 20 scrambled control images which should be passively viewed. Each image was shown on the screen for a fixed duration of six seconds. To maintain participants’ attention, ten trials were randomly chosen, in which participants had to indicate (via keypress) whether the last presented image depicted a person or not. We included an intertrial-interval (ITI) between consecutive image presentations, which was marked by a white fixation cross. The duration of the ITI was sampled from a poisson distribution (M = 2 s; range: 1-9 s). The total duration of the cue phase was ten minutes. Following the first cue phase, participants completed 128 trials of a classical delay-discounting task (Peters & Büchel, 2009). On each trial, participants chose between a fixed immediate reward of 20€ (SS) and a variable delayed amount (LL). Every trial started with the presentation of the available LL-reward and the associated delay (e.g., 38€, 14 days). The LL-reward was depicted centrally on the screen for a fixed duration of two seconds. LL-presentation was followed by a short jitter interval which was marked by a white fixation cross. The duration of the jitter interval was sampled from a poisson distribution (M = 2 s; range: 1-9 s) and was followed by the decision screen. Here, participants chose between one of two symbols corresponding to the two options (SS: circle; LL: square). The response window was four seconds. The chosen option was highlighted for one second. The ITI was again marked by a white fixation cross with a presentation duration sampled from a poisson distribution (M = 2 s; range: 1-9 s). An example trial is depicted in **Figure 1**.

**Figure 1.**
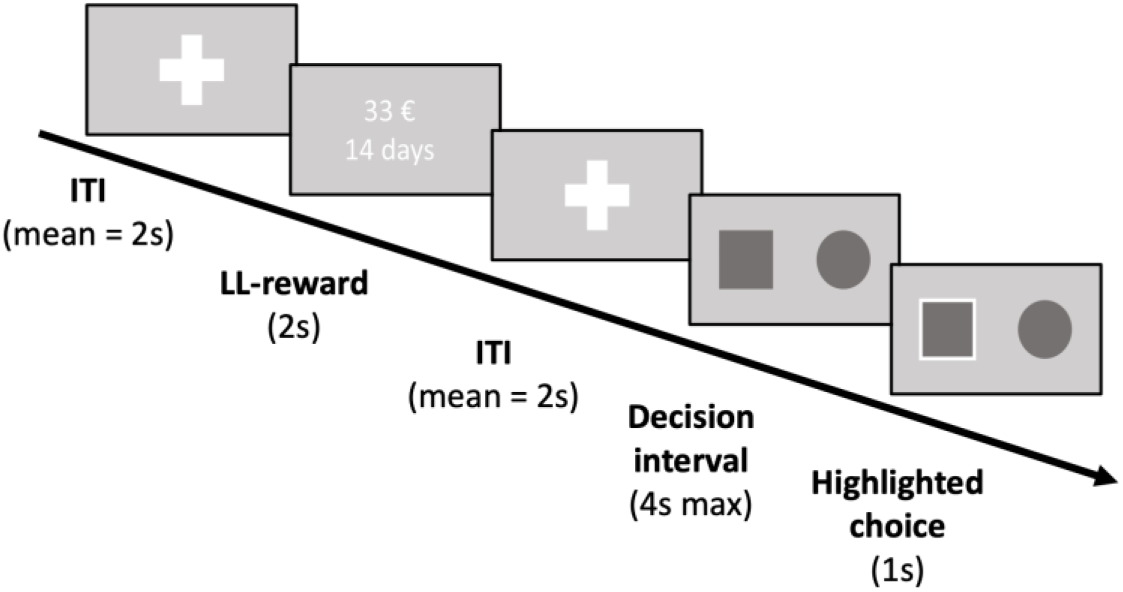
Example trial from the delay discounting task. Larger later reward (LL) presentation was preceded and followed by short jitter intervals (ITI), marked by white fixation crosses; Durations for the jitter intervals were sampled from a poisson distribution (M = 2 s; range: 1-9 s); Thereafter, the decision screen was presented. The fixed smaller sooner reward (SS; 20€) was never shown throughout the experiment.

The LL-rewards were calculated beforehand by multiplying the SS-amount with two different sets of multipliers, differing slightly across days (Set 1: [1.01 1.02 1.05 1.10 1.15 1.25 1.35 1.45 1.65 1.85 2.05 2.25 2.65 3.05 3.45 3.85]; Set 2: [1.01 1.03 1.08 1.12 1.20 1.30 1.40 1.50 1.60 1.80 2.00 2.20 2.60 3.00 3.40 3.80]). We likewise used two sets of delays (Set 1: [1 3 5 8 14 30 60 122 days]; Set 2: [2 4 6 9 15 32 58 119 days]). Multiplier and delay combinations were randomly assigned to testing days per participant. Participants were instructed explicitly about the task structure and performed ten test trials during a practice run within the scanner. In accordance with previous studies (Green et al., 1997; Mathar et al., 2022; Wagner et al., 2020) we used hypothetical choice options. However, note that discount rates for real and hypothetical rewards are highly correlated and similarly processed on the neuronal level (Bickel et al., 2009; Johnson & Bickel, 2002).

Following the TD task, participants underwent a second analogous cue phase, which was then followed by a reinforcement learning task (Two-Step task). This task is preregistered separately and will be reported elsewhere.

The second day followed exactly the same structure, with the exception of the cue phases. Depending on the condition on the first day, participants were presented with images from the other condition (neutral or erotic). The sequence was counterbalanced between participants (50% of the participants started with the erotic cue condition, the other 50% were first presented with neutral cues). After completing the scanning session on the second day, participants performed three short working memory tasks (operation span (Foster et al., 2015), listening span (van den Noort et al., 2008), and digit span (Wechsler, 2008)) on a laptop and filled out a computerized questionnaire battery as well as a demographic survey. However, note that data from demographic, health and personality questionnaires will be reported elsewhere.

### 2.4. Data analysis

#### 2.4.1. Behavioral data analysis of intertemporal choice

We used two different approaches to quantify impulsivity as measured by the TD task. Our model-based approach assumed hyperbolic devaluation of delayed rewards (Green & Myerson, 2004; Mazur, 1987) and a softmax choice rule for modeling subjects’ intertemporal decisions. For model-free analysis we directly focused on actual choice preferences of SS- and LL-options.

##### Model-agnostic approach

A model-free analysis can avoid problems associated with the choice for a particular theoretical framework (e.g., hyperbolic discounting) or extreme parameter estimates that result in skewed distributions. The latter might yield problems for statistical approaches that require normally distributed variables. We therefore simply computed the relative proportion of SS-choices for every participant and condition (neutral vs. erotic) to obtain a model-agnostic measure of TD (Eq. 1).

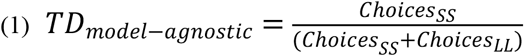

##### Model-based analysis

We used hierarchical Bayesian modeling to fit a hyperbolic discounting model with softmax action selection to the choice data. For each parameter (discount rate *k*, modelled in log-space, and inverse temperature *ß*) we fit separate group-level Gaussian distributions for the neutral condition from which individual subject parameters were drawn. To model condition effects on each parameter, we fit separate group-level distributions modeling deviations from the neutral condition for erotic cues, respectively (“shift”-parameter; Eqs. 2 -3).

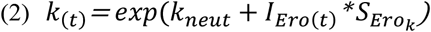

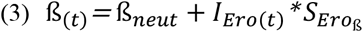

Here, *I*_*Ero*_ is a dummy-coded indicator variable coding the experimental condition (1=erotic, 0=neutral) and *S*_*Ero*_ are subject-specific parameters modeling changes in log(*k*) and *ß* depending on the condition in trial *t*. Computation of the discounted subjective value (*SV*) of the larger later option (*LL*) for a given delay *D* and amount *A* in a given trial then uses the standard hyperbolic model (Eq. 4):

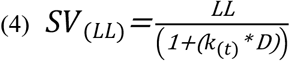

However, cue exposure might also affect TD beyond a modulation of log(*k*), e.g., by inducing an overall offset in the discounting function. To account for such effects, we examined another model that allowed for an offset in the discounting function in the neutral condition (modelled by the parameter 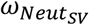), which might then again be differentially affected by erotic cues (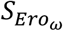, Eq. 5).

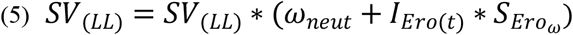

Because a positive 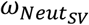 would indicate a subjective value of the LL that exceeds the objective amount (at delay = 0), the range of the offset-parameter was restricted between 0 and 1. Finally, subjective values of SS- and LL-options as well as modulated inverse temperature parameter *ß* (Eq. 3) were then used to calculate trial-wise choice probabilities according to a softmax choice rule:

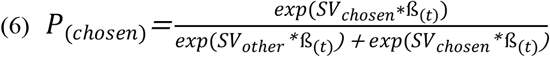

In summary, we compared two models: Model 1 (*Base-model*) only included 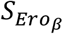 and 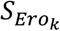 to assess cue exposure effects on ß and log(k). Model 2 (*Offset-model*) additionally included a potential change in the offset, 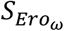.

##### Parameter estimation

Posterior parameter distributions were estimated via no-U-turn sampling (NUTS; Hoffmann & Gelman, 2014) implemented in STAN (Carpenter et al., 2017) using R (R Core Team, 2018) and the RStan Package (Stan Development Team and others, 2018). Prior distributions for the group-level parameters are listed in **Table 1**. Group mean priors were derived from posterior means and standard deviations from a recent study from our group, based on the Base-model (Mathar et al., 2022). STAN model code for all models is publicly available at OSF (Base-Model: osf.io/6uz8g; Offset-Model: osf.io/mgjx5). Model convergence was assessed via the Gelman-Rubinstein convergence diagnostic 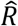 and values of 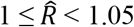 were considered acceptable. We ran 4 chains with a burn-in period of 1500 samples and no thinning. 4000 samples were then retained for further analysis. For details on MCMC convergence, see Gelman and Rubin (1992). We used Bayesian statistics (see Kruschke, 2010) to evaluate cue effects on model parameters of the best fitting model. Relative model fit was assessed via the loo-package in R using the Widely-Applicable Information Criterion (WAIC), where lower values reflect a superior fit of the model while considering its complexity (Vehtari et al., 2017; Watanabe, 2010).

**Table 1.** Priors of group-level parameter means

| Base-model | Parameter | Group mean prior |
| --- | --- | --- |
| | $k$ | Normal (-4.2, 2.01) |
| | $S_{Ero_k}$ | Normal (.15, .64) |
| | $\beta$ | Normal (.51, .3) |
| | $S_{Ero_\beta}$ | Normal (.02, .11) |
| Offset-model | Parameter | Group mean prior |
| | $k$ | Normal (-4.2, 2.01) |
| | $S_{Ero_k}$ | Normal (.15, .64) |
| | $\beta$ | Normal (.51, .3) |
| | $S_{Ero_\beta}$ | Normal (.02, .11) |
| | $\omega$ | Uniform (0, 1) |
| | $S_{Ero_\omega}$ | Normal (0, .4) |

We analyzed posterior distributions of group mean condition effects (as reflected in the *S*_Ero_ parameters, see Equations 2, 3 and 5 above) by computing their highest density intervals (HDI) and assessed their overlap with zero. We further report *undirected* Bayes factors (BF01) based on the Savage-Dickey-Density Ratio, which quantify the degree of evidence for a null model that would restrict a parameter of interest at a given value (e.g., *S*_*Ero*_ = 0) against an alternative model where the parameter can vary freely (see Marsman & Wagenmakers, 2017 for details). To test the degree of evidence for increases vs. decreases in parameter values, we computed *directional* Bayes factors (dBFs) for condition effects. A dBF corresponds to the ratio of the posterior mass of the shift-parameter distribution below zero to the posterior mass above zero (Marsman & Wagenmakers, 2017). We considered Bayes Factors between 1 and 3 as anecdotal evidence, Bayes Factors above 3 as moderate evidence and Bayes Factors above 10 as strong evidence. Likewise, the inverse of these values reflect evidence in favor of the opposite hypothesis (Beard et al., 2016).

##### Posterior predictive checks

We used posterior predictive checks to assess the degree to which the included models (Base-model, Offset-model) reproduced key patterns in the data, in particular the change in LL choice proportions across delays. For this purpose, we simulated 4000 datasets from each model’s posterior distribution and plotted the mean observed proportion of LL choices and the simulated LL choice proportions across delay. This was done separately for both conditions (neutral, erotic).

#### 2.4.2. fMRI data acquisition

MRI images were acquired on a 3 Tesla Magnetom Prisma Fit system (Siemens, Erlangen, Germany) equipped with a 64-channel head coil. Task stimuli were presented on an MR compatible screen and rearview mirror system. Participants responded with their index and middle fingers on a two-button box, held in their right hand. Psychophysics Toolbox Version 3.52 implemented within MATLAB R2019b software (The Mathworks Inc., MA, USA) was used for stimulus presentation and behavioral data collection. Functional images were acquired in 5 separate runs (Cue phase1, TD, Cue phase 2, Two-step (first half), Two-step (second half)) by utilizing a multiband gradient echo-planar imaging (mb-EPI) sequence with repetition time (TR) = 0.7 s, echo time (TE) = 37 ms, flip angle = 52°, field of view (FOV) = 208 mm, voxel size = 2 mm^3^ isotropic (slice thickness = 2 mm, no gap), and multiband acceleration factor of 8. Each volume consisted of 72 transverse slices acquired in alternating order from the anterior-posterior commissure plane. The 5 runs contained ∼7700 volumes for each participant and ∼90 min. of total scan time per day.

#### 2.4.3. fMRI data analysis

Preprocessing and analyses of fMRI data was performed using SPM12 (v7771; Wellcome Trust Centre for Neuroimaging) implemented in MATLAB R2019b (The MathWorks), and the FMRIB Software Library (FSL; Version 6.0.4; Jenkinson et al., 2012). Prior to statistical analysis, the first five functional volumes were discarded to allow for magnetic saturation. Functional images were corrected for motion and distortion artifacts using the FSL tools MCFLIRT and topup (Andersson et al., 2003; Smith et al., 2004). Next, anatomical T1-images were co-registered to functional images and normalized to the Montreal Neurological Institute (MNI) reference space using the unified segmentation approach in SPM12. Finally, we normalized functional images using the ensuing deformation parameters, and smoothed using a 6mm full-width-half-maximum Gaussian kernel.

##### Cue phase

*1*^*st*^*/2*^*nd*^ *level modeling*. Both testing days entailed two separate cue exposure phases (session 1 & 2). Note that to examine cue effects on TD, we only focused on the first cue exposure session directly preceding the TD task. In each cue phase, participants viewed 40 intact and 20 scrambled images. Depending on the condition of the day, intact images depicted either everyday scenes and portraits of people (neutral condition) or nude women (erotic condition).

Using SPM12, functional images from each day were analyzed using a general linear model (GLM). Each GLM examined the sustained activity during the presentation of intact and scrambled image types using boxcar regressors (duration = 6s) which were convolved with the canonical hemodynamic response function (HRF). To account for residual variance caused by subject movement we included the following nuisance regressors: 24 motion parameters (6 motion parameters relating to the current and the preceding volume, respectively, plus each of these matrices squared, see Friston et al., 1996), mean signal extracted from the ventricular cerebrospinal fluid (CSF), and a matrix containing motion-outlier volumes (identified by assessing global intensity differences between subsequent volumes; threshold: > 75th percentile + 2.5 * interquartile range of the global signal).

Contrast images for intact and scrambled image presentation from the cue exposure phases (Cuephase1_Erotic;_ Cuephase1_Neutral_) were then entered into a second-level random effects model (flexible factorial design; factors: subjects, visibility (intact, scrambled), condition (erotic, neutral)) to evaluate BOLD-activity changes attributable to erotic image content. Variances for all factors were set to: *equal*. We included a main effect for “subject” and an interaction term for the factors “visibility” and “condition”.

We ran two analyses to evaluate neural effects of neutral vs. erotic cues. First, to replicate erotic cue effects (vs. intact neutral cues), we examined a priori-defined regions-of-interest (ROIs) related to the processing of visual erotic stimuli (see H4; Stark et al., 2019). The ROI mask was created using the group-level results (t-map) for the contrast erotic > neutral from Stark et al. (2019). We applied a binarization threshold (t-value = 6) to extract voxels showing increased responsiveness to visual sexual stimuli, and then used small volume family wise error (FWE) correction (*p*<0.05) across the entire mask to control for multiple comparisons. Further whole-brain effects of visual cue exposure are reported at a FWE corrected threshold (*p* < 0.05; peak-level).

Second, we tested for associations between reward-system activity (erotic>neutral) within key dopaminergic (Nacc, VTA) and prefrontal (DLPFC) regions and behavioral cue effects on TD following erotic vs. neutral cue exposure (see H6). In more detail, we assessed associations between neuronal cue-reactivity-responses within the first cue phase (Erotic_session1_ > Neutral_session1_) and subject-specific shift-parameters (S_Ero(k),_ S_Ero(ω)_), capturing condition-specific changes in TD. Associations were quantified via Bayesian correlations (using JASP (JASP Team, 2022; Version 0.14.3)) separately for pre-defined subcortical (Nacc, VTA) and cortical (DLPFC) ROIs.

To extract VTA activity, we first constructed an anatomical mask based on Stark and colleagues (2019; see above). Specifically, we used reported peak coordinates from the group contrast erotic>neutral (VTA: -6, -8, -10 and the mirrored location) as centers of two 10mm spherical ROIs, which we then combined into a bilateral mask. For Nacc, we focused on the striatal cluster within the “reward” mask based on two metanalyses, provided by the Rangel Neuroeconomics lab (http://www.rnl.caltech.edu/resources/index.html). This mask combines bilateral striatum, vmPFC, posterior cingulate cortex (PCC) and anterior cingulate cortex (ACC). Lastly for DLPFC, we built a custom mask based on previous studies reporting increased DLPFC-activity during LL vs. SS choices (Smith et al., 2018; see below). To calculate brain-behavior correlations, we first identified peak voxels from our group-level contrast erotic(intact)>neutral(intact) within the mentioned VTA, striatum and DLPFC masks and extracted parameter estimates from these voxels for each participant.

##### Delay-discounting-task

*1*^*st*^*/2*^*nd*^ *level modeling*. On both testing days, the first cue exposure phase was followed by a classical delay discounting task (see methods section 2.3). Functional images from both days (i.e., conditions) were analyzed separately using general linear models (GLM) implemented in SPM12. Each GLM included the following regressors: (1) the presentation of the larger later option (LL) as event regressor (duration = 2s), standardized discounted subjective value (SV) as parametric modulator (computed based on the best-fitting model), (3) the onset of the decision period as stick regressor (duration = 0s) and (4) the choice (LL vs. SS) as parametric modulator. Invalid trials on which the participant failed to respond within the response window (limit: 4s) were modeled separately. All regressors were convolved with the canonical hemodynamic response function as provided by SPM12. Residual movement artifacts were corrected by using the same nuisance regressors as for the cue phase (see above).

We hypothesized subjective value (SV) of delayed rewards to be encoded in ventral striatal (VS) and ventromedial prefrontal areas (vmPFC) and that lateral prefrontal cortex activity (DLPFC) would be increased during choices of LL rewards (see H2 & H3). Further, we predicted that DLPFC activity during delayed reward presentation would be reduced following erotic cue exposure (see H5). To test H2 and H3, we entered the respective contrast images of parametric effects of subjective value (SV) and the chosen option (LL vs. SS) into separate second-level random effects models. We focused on pre-defined ROIs implicated in TD SV-computations (H2; Bartra et al., 2013; Clithero and Rangel, 2014) and choice behavior (H3; Smith et al., 2018). Specifically, H2 was tested using again the combined “reward” mask, which was provided by the Rangel Neuroeconomics lab (http://www.rnl.caltech.edu/resources/index.html), and combines bilateral striatum, vmPFC, PCC and ACC. To test H3 we again used the above-mentioned DLPFC-mask created on findings on LL vs. SS choices (Smith et al., 2018). To control for multiple comparisons, we applied small volume correction (SVC; *p*<0.05, peak-level) across the reward mask (H2) or the DLPFC mask (H3).

Finally, we tested for condition-related (erotic vs. neutral) differences in prefrontal activation related to the onset of the LL-rewards (H5). For this purpose, LL-onset regressors were directly compared between neutral and erotic image conditions on the group level. Here we again used the above mentioned preregistered DLPFC-ROI (Smith et al., 2018) for SVC (*p*<0.05, peak-level).

#### 2.4.4. Deviations from preregistered analyses

This study was preregistered (https://osf.io/w5puk). We deviated from the pre-registered analyses in the following ways: First, based on mean effect size estimates from two previous studies on erotic cue exposure effects on TD, we preregistered a minimum sample size of n=31 to reach a power of .80 (effect size Cohen’s f = 0.22, error probability α = .05). To account for potential dropout, we aimed for a final sample size of n=40. Due to technical issues of the MRI scanning environment and the final sample consisted of 38 subjects which was further reduced to 36, as two participants voluntarily dropped out of the experiment. Nevertheless, this still exceeds minimum sample size by 5, indicating we had enough power to detect potential erotic cue effects on TD.

Second, we slightly deviated from our planned computational modeling approach to quantify erotic cue effects on TD. We initially preregistered three models which all used hierarchical Bayesian modeling to fit variants of the hyperbolic model with softmax action selection to the choice data. However, two of the preregistered models suffered from two shortcomings (Model 2 & 3 in the preregistration). First, they both assumed cue-induced SV-offsets *only in the erotic condition*, thereby selectively increasing flexibility and predictive power in one condition. To correct this asymmetry, we now allowed for an offset of the discounting function in the neutral condition, which again could be differentially modulated by erotic cues (see Model 2, section 2.4.1). Second, the preregistered offset-parameter was initially defined as additive. However, validation analyses revealed that this formulation yielded implausible SVs (e.g., SV_LL_ < 0) in a few individuals that exhibited extremely unbalanced choice behavior (e.g., only very few SS or LL choices). Therefore, we changed the model formulation to a multiplicative offset (see Eq. 5).

#### 2.4.5. Data and code availability

T-maps of 2^nd^ level contrasts as well as STAN model code and raw behavioral data are available on the Open Science Framework (T-maps: https://osf.io/9uzm8/; Stan model code: Base-Model: osf.io/6uz8g; Offset-Model: osf.io/mgjx5; Raw data: https://osf.io/nxcas).

## 3. Results

The results section is structured as follows. In accordance with our preregistered analysis plan, we first report the results of the replication analyses for the fMRI data for subjective value coding (H2), intertemporal choice (H3) and erotic cue processing (H4). Next, we report behavioral and modeling results regarding effects of cue exposure on TD (H1). Finally, to link neuronal cue-reactivity to TD, we report findings from two separate analyses. First, we assessed cue exposure effects on DLPFC activity at the during LL-reward presentation (H5). Second, we examined between-subjects associations between erotic reward-system-responsivity within key dopaminergic (Nacc, VTA) and prefrontal (DLPFC) areas, and alterations in TD (H6).

### 3.1. Neuronal correlates of subjective discounted value

We hypothesized subjective value (SV) coding of delayed rewards in striatal (VS) and ventromedial prefrontal areas (vmPFC; see Peters & Büchel, 2009; see H2). Our GLM incorporated the onset of the LL-option as event regressor (duration = 2s) and the standardized discounted subjective value (SV) of the LL option as parametric modulator. SVs were based on the best-fitting *Offset-model* (methods section 2.4.2, lowest WAIC, see below). We used a combined “reward” ROI mask provided by the Rangel Neuroeconomics lab (http://www.rnl.caltech.edu/resources/index.html). This mask combines bilateral striatum, vmPFC, PCC and ACC and was used for small-volume correction.

Figure 2A shows brain activation that covaried with subjective discounted value of larger later rewards across experimental conditions (main effect across *erotic* and *neutral*). ROI analysis replicated previous findings on subjective value coding in a large cluster within medial prefrontal cortex (peak coordinates x, y, z (in mm): 2, 54, -10; *z*-value = 5.83, *p*_SVC_ < 0.001), posterior cingulate cortex (PCC; - 10, -34, 38; z-value = 4.38, *p*_SVC_ = 0.005) and right ventral striatum/caudate (VS; 4, 10, 2; z-value = 4.22, *p*_SVC_ = 0.010) – confirming hypothesis H2. Parameter estimates extracted from vmPFC, PCC and VS peak voxels illustrate that this effect was evident in the vast majority of individual participants (see Figure 2B). Value-related activity in predefined ROIs did not differ between experimental conditions (no suprathreshold clusters for the contrasts: erotic > neutral or neutral > erotic). When running separate analyses for each condition, significant SV coding was confirmed in mPFC, PCC and VS in the erotic condition (*p*_SVC_ < 0.05). In the neutral condition, this was true for the mPFC, VS reached trend level (see **Supplementary Table S1**). T-maps from the respective group-level contrasts are publicly available at OSF (https://osf.io/9uzm8/).

**Figure 2.**
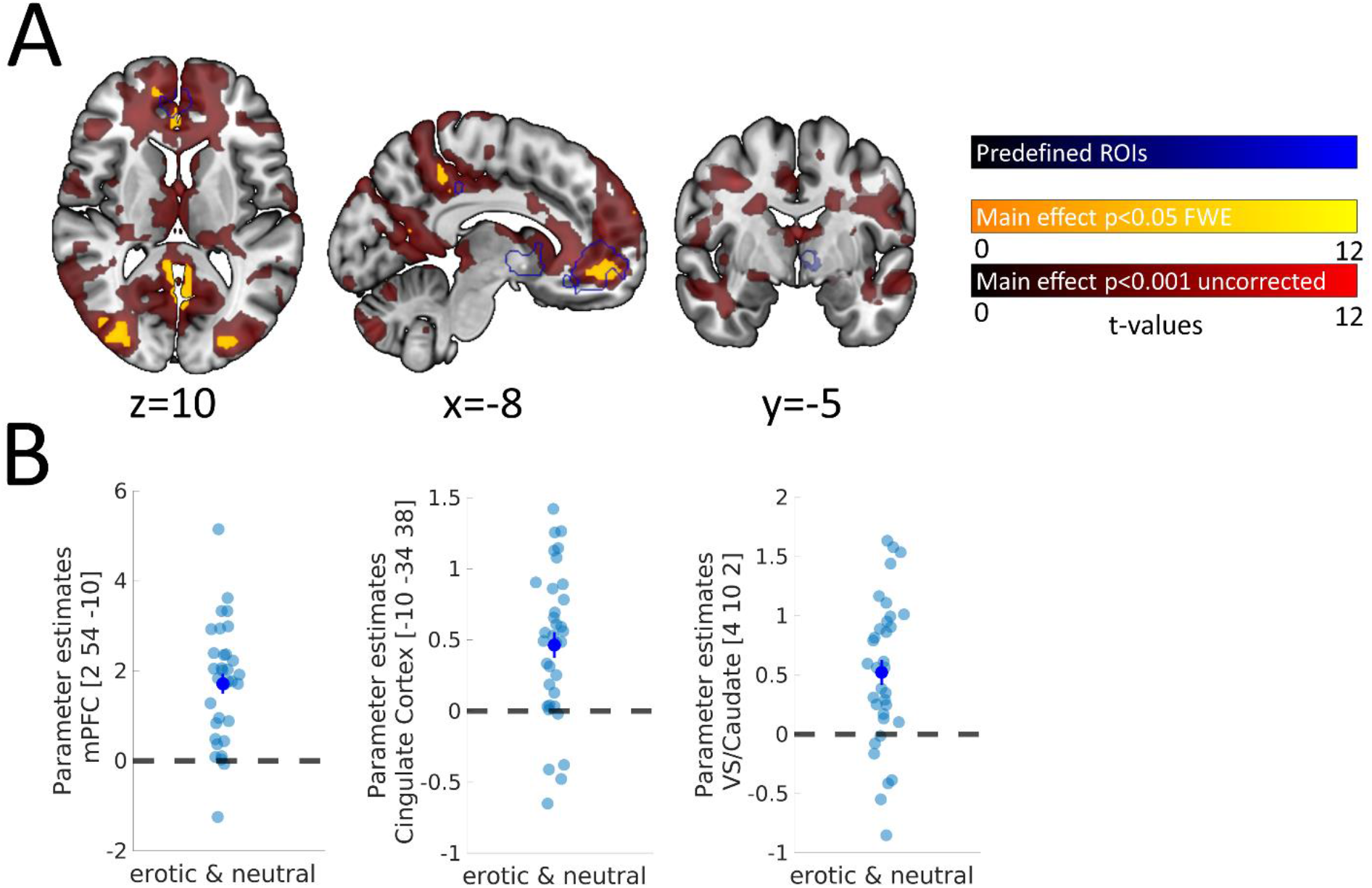
Neuronal correlates of subjective value (SV). **A**: Display of the parametric SV-regressor (main effect across conditions); red, *p* < 0.001 (uncorrected); yellow, whole-brain FWE corrected *p* < 0.05; blue, preregistered regions of interest from reward mask (see above); **B**: Extracted *ß*-estimates of each participant extracted from medial prefrontal cortex (mPFC), posterior cingulate cortex (PCC) and ventral striatum/caudate (VS) peak coordinates of the parametric SV-regressor; Error bars denote SEM.

### 3.2. Neuronal correlates of intertemporal choice

We predicted increased dorsolateral prefrontal cortex activity (DLPFC) during choices of LL vs. SS rewards (Smith et al., 2018; see H3). Our GLM included the onset of the decision period as event regressor (duration = 0s) and the selected option (LL vs. SS) as parametric modulator. We built (and preregistered) a custom (left) DLPFC-mask based using a 12 mm sphere centered at the group peak coordinates for the contrast ‘LL-vs. SS-choice’ reported by Smith and colleagues (2018) (peak coordinates (x = -38, y = 38, z = 6).

This ROI analysis replicated increased activity in left DLPFC associated with LL vs. SS choices across conditions (main effect across *erotic* and *neutral;* peak coordinates: -40, 48, 4; z-value = 4.26, *p*_SVC_ = 0.003; see **Figure 3**), confirming hypothesis H3. We found no suprathreshold clusters for the contrasts: erotic > neutral or neutral > erotic. This effect was also confirmed in our preregistered ROI when each condition was analyzed separately (**Supplementary Table S2**). T-maps from the respective group-level contrasts are publicly available at OSF (https://osf.io/9uzm8/).

**Figure 3.**
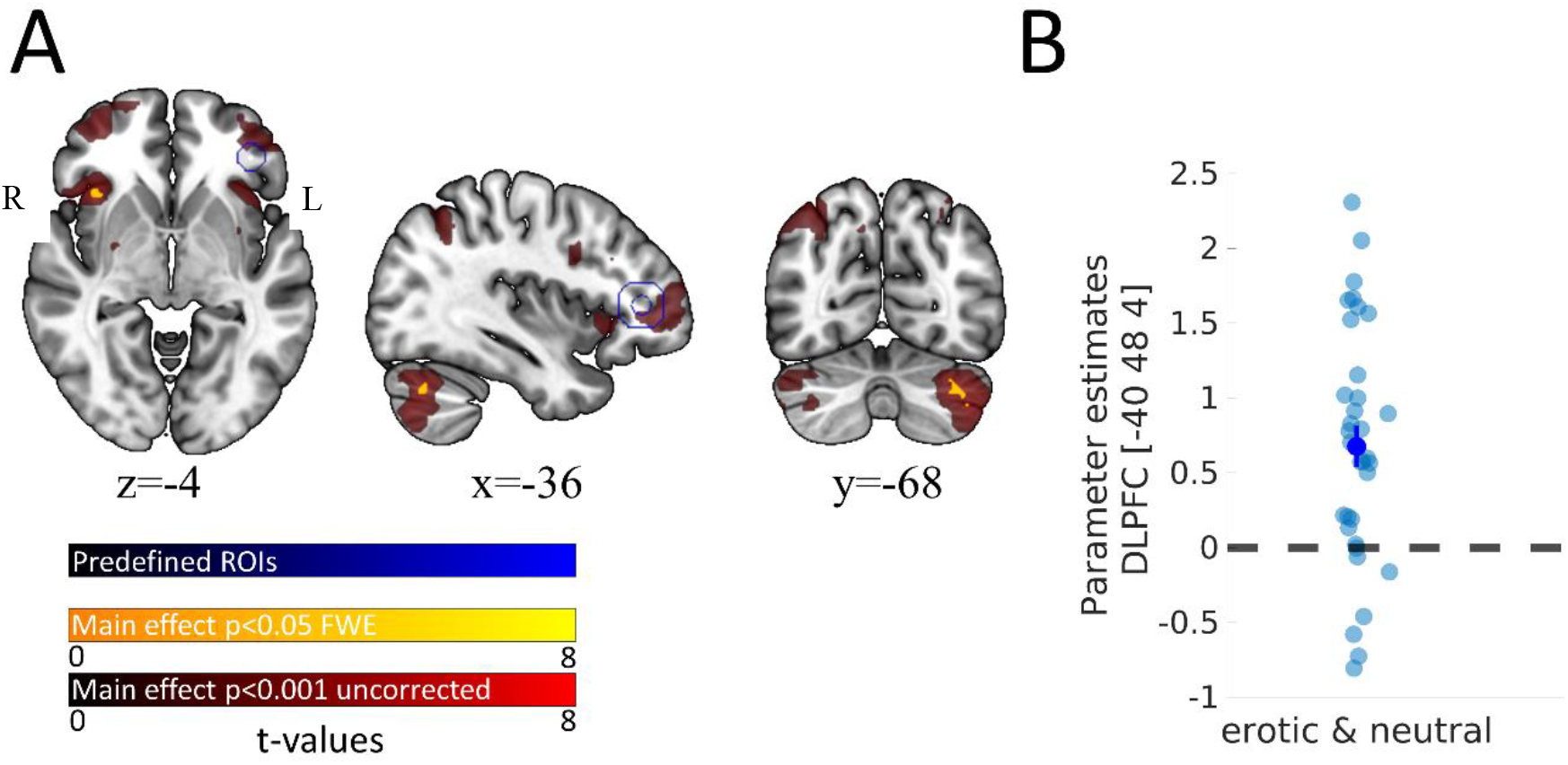
Neuronal correlates of larger-later (LL) vs. smaller-sooner (SS) choices. **A**: LL > SS contrast (main effect across conditions); red, *p* < 0.001 (uncorrected); yellow, whole-brain FWE corrected *p* < 0.05; blue, predefined regions of interest from custom DLPFC mask (see above); **B**: *ß*-estimates of each participant extracted from left DLPFC peak coordinates; Error bars denote SEM.

Subsequent whole brain (FWE-corrected) analysis revealed two additional clusters coding for LL vs. SS choices across conditions (main effect across *erotic* and *neutral)*, located in the right insular cortex (36, 20, -4; z-value = 5.38, *p*_FWE_ = 0.007) and the cerebellum (−34, 66, -34, z-value = 5.28, *p*_FWE_ = 0.012). We found no suprathreshold clusters for either condition contrast (erotic > neutral; neutral > erotic) using whole-brain FWE correction (*p*<0.05).

In an exploratory whole brain approach, we also checked for brain activity associated with choices of the immediately available option, that is the smaller but sooner option/reward (SS). Here we found that brain activity within a multitude of cortical (cerebellum, mid cingulate, bilateral insula, mid frontal cortex) but especially subcortical regions (bilateral caudate, right putamen, thalamus, hippocampus) positively correlated with SS-choices across both experimental conditions (see **Supplementary Figure S1**). For the condition contrasts, erotic > neutral and neutral > erotic however, no voxels survived whole-brain FWE correction (*p*<.05).

### 3.3. Appetitive cue effects on neuronal reward circuitry

We predicted (erotic-) cue effects on TD to be at least partly moderated by activations in neuronal reward circuits (Li, 2008; Stark et al., 2019; Yeomans & Brace, 2015). During the cue exposure phase, participants were exposed to 40 intact (erotic or neutral) and 20 scrambled control images. Analyses only focused on the first cue exposure session directly preceding the TD task. We ran a flexible factorial random-effects model (factors: visibility (intact/scrambled), condition (erotic/neutral)) and preregistered ROIs based on a previous study (Stark et al., 2019; see methods section for details). ROI analyses applied small volume FWE correction (*p*<0.05) across the entire mask.

A sanity check confirmed widespread functional responses across occipital and ventral temporal cortices for the intact vs. scrambled contrast (see **Supplementary Figure S2**).

As depicted in **Figure 4**, (intact) erotic, compared to (intact) neutral cue exposure was associated with increased activity in widespread cortical and subcortical regions. Our preregistered ROI analysis revealed increased activity in four large posterior (cortical) clusters for erotic vs. neutral cues, including right inferior temporal cortex (52, -60, -4; z value = 6.25, *p*_SVC_ < 0.001), left inferior occipital cortex (−48, -68, -6; z value = 5.42, *p*_SVC_ = 0.001), right superior parietal cortex (26, -60, 62, z-value = 4.76, *p*_SVC_ = 0.013) and right middle occipital cortex (28, -72, 30; z value = 4.51, *p*_SVC_ = 0.036).

**Figure 4.**
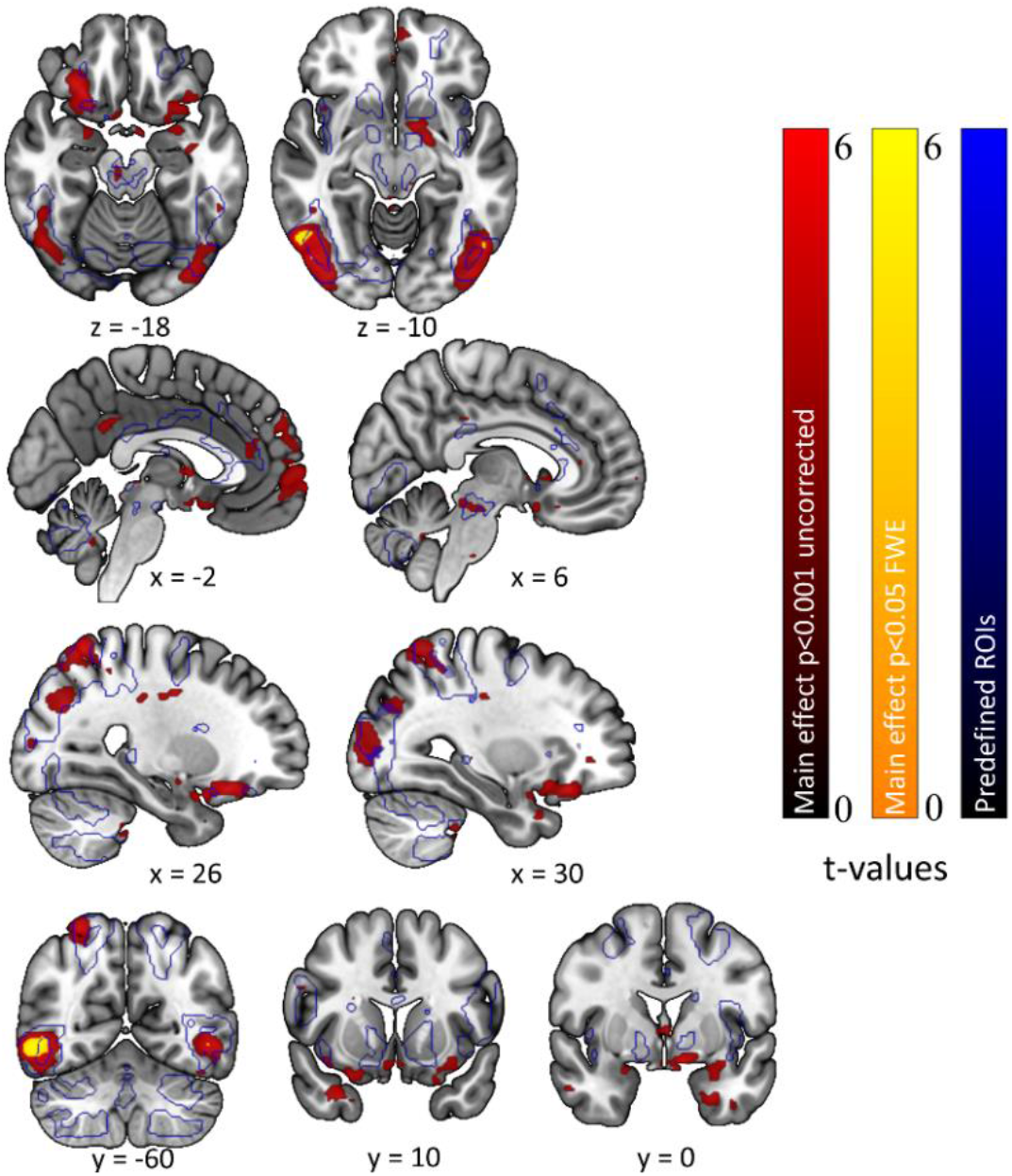
Neuronal correlates of (intact) experimental image processing (erotic > neutral). Red, *p* < 0.001 (uncorrected); yellow, whole-brain FWE corrected *p* < 0.05; blue, predefined regions of interest from ROI mask (see above).

We had predicted subcortical activations in reward-related brain regions (e.g., VS, vmPFC) to be linked to erotic cue exposure (H4), but many subcortical effects fell just beyond the preregistered ROI-mask based on Stark et al. (2019). We therefore followed up with a second (not preregistered) ROI analysis using the above-mentioned “reward” mask, based on two meta-analyses, provided by the Rangel Neuroeconomics Lab (http://www.rnl.caltech.edu/resources/index.html). Small volume correction was again applied across the entire mask. As expected, this confirmed highly robust bilateral effects in the VS/caudate (left: -10, 2, -10; z-value = 4.59; *p*_SVC_ = 0.002; right: 4, 6, 2; z-value = 3.70; *p*_SVC_ = 0.047) and the vmPFC (−6, 58, -2; z-value = 4.54; *p*_SVC_ = 0.002; see **Figure 5**).

**Figure 5.**
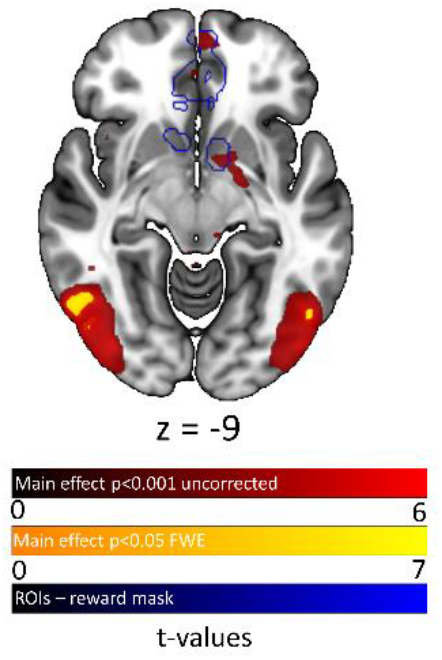
Neuronal correlates of (intact) experimental image processing (erotic > neutral). Red, *p* < 0.001 (uncorrected); yellow, whole-brain FWE corrected *p* < 0.05; blue, regions of interest from reward mask (not preregistered, see above).

A t-map depicting all activations associated with erotic > neutral image processing is publicly available at OSF (https://osf.io/9uzm8/). We also checked for increased brain activity following neutral compared to erotic image presentation. However, here we identified no suprathreshold clusters.

### 3.4. Appetitive cue effects on intertemporal choice

Having thus replicated previous findings on subjective value coding (H2), intertemporal choice (H3) and erotic stimulus processing (H4) (Peters & Büchel, 2009; Smith et al., 2018; Stark et al., 2019), we next assessed condition-related changes in TD behavior.

#### Model-agnostic analysis

Contrary to our hypothesis (H1), TD was not differentially affected by appetitive cue exposure (see **Figure 6**). In the neutral condition, the SS-option was chosen in 39.6% of trials whereas in the erotic condition the SS-option was chosen in 38.5% of trials (t_(35)_ = 0.714, *p* = 0.480).

**Figure 6.**
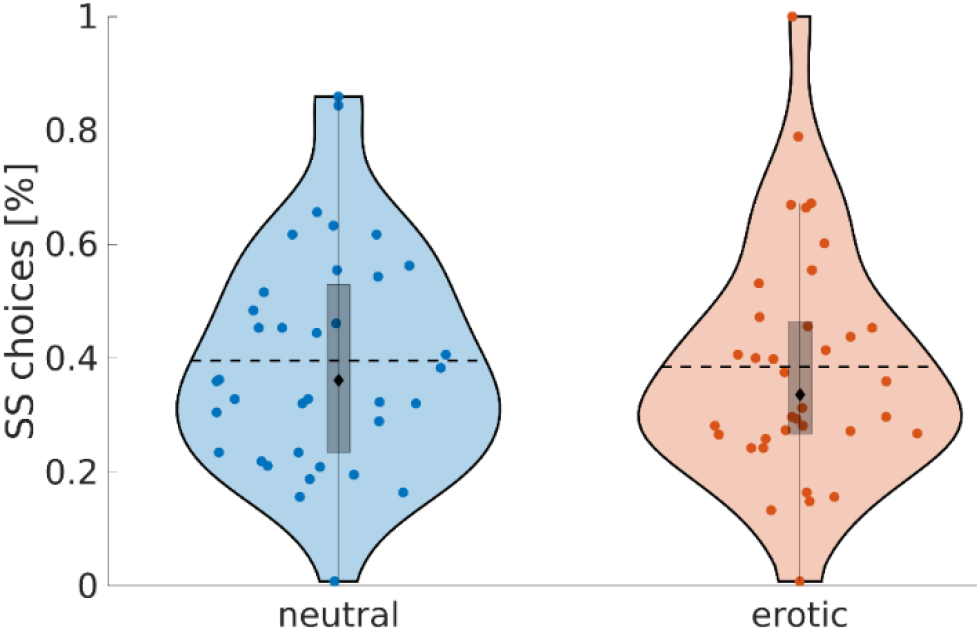
Percentage of smaller-sooner choices split by experimental condition (neutral vs. erotic). Colored dots = single subjects; Dashed lines = condition means; Black diamonds = condition medians; The edges of the boxes depict the 25th and 75th percentiles, the whiskers extend to the most extreme datapoints the algorithm considers to be not outliers.

#### Computational modeling

Model comparison revealed, that choice data were best captured by a hyperbolic model with an additional SV-offset-parameter *ω*, in addition to parameters accounting for choice consistency (*ß*) and steepness of TD (log(*k*); Offset-model). This model comparison replicated across conditions (neutral, erotic), and was confirmed in the combined model including parameters modeling condition effects (see **Table 2**). The superior fit of the offset-model was also reflected in choice predictions. The Offset-model accounted for around 82.2% (Base-model: 79.6%) of all decisions (**Supplementary Table S3, Supplementary Figure S3**). Finally, posterior predictive checks confirmed that LL-choice proportions across delays were much better accounted for by the Offset-model (**Figure 7**). All further analyses therefore focused on the Offset-model. However, note that due to an extreme behavioral choice pattern (only one single SS-choice in both conditions), data from one participant could not be explained by our winning model and was excluded from all further analyses.

**Table 2.** Summary of the WAICs of all included hyperbolic models in all sessions. Ranks are based on the lowest WAIC.

| Model | Neutral Condition |  | Erotic Condition |  | Combined |  |
| --- | --- | --- | --- | --- | --- | --- |
|  | WAIC | Rank | WAIC | Rank | WAIC | Rank |
| Base-model | 2654.537 | 2 | 2699.152 | 2 | 5365.867 | 2 |
| Offset-model | 2292.945 | 1 | 2453.293 | 1 | 4771.835 | 1 |

Note. WAIC = Widely applicable information criterion

**Figure 7.**
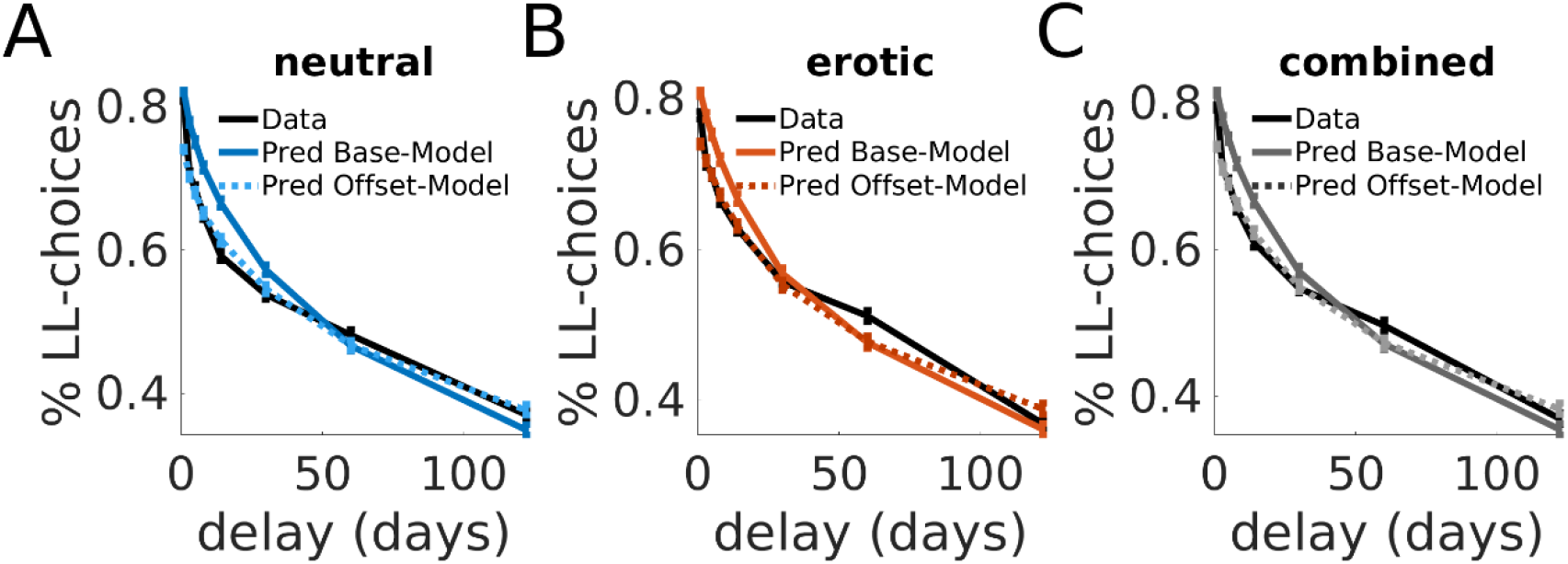
Group level posterior predictive checks for the included temporal discounting models (Base-model, Offset-model). Here we plotted the mean observed proportion of LL-choices and the simulated LL-choices from both models for each delay. Specifically, we created 4k simulated data sets from each model’s posterior distribution. For each simulated participant, we calculated the fraction of LL-choices across eight delay bins. Next, we calculated group average proportion of LL-choices for each delay and associated standard errors (vertical bars). Simulated data were then overlaid over the observed choice data. We did this separately for the neutral (A) and erotic (B) conditions as well as for the combined datasets (C).

Examination of the posterior distributions of the best-fitting model then confirmed the model-agnostic results. TD (log(*k*); **Figure 8A**) was not substantially affected by erotic cue exposure (SERO_(k)_; **Figure 8B**), such that the highest density intervals for SERO_(k)_ substantially overlapped with zero. These data were more likely to be observed under a null hypothesis assuming SERO_(k)_ to be equal to zero (BF01 = 4.11). Interestingly, SV-offset parameters *ω*_neutral_ clearly differed from one in all participants, emphasizing the general utility of this additional parameter to account for a choice bias irrespective of delay. However, the observed data were much more compatible with the null model where the condition effect in the offset was equal to zero (BF01 = 43.18; **Figure 8 C, D**), strongly suggesting the offset was not modulated by erotic cue exposure. Likewise, data for the SERO_(*ß*)_ parameter were much more compatible with the null model, indicating the change in stochasticity following erotic cue exposure was equal to zero (BF01 = 18.473, see **Figure 9 A, B**). See **Table 3** for summary statistics and Bayes factors of the posterior distributions of all relevant parameters. For completeness, posterior distributions and Bayes factors from the inferior Base-model are reported in the supplement **(Figure S4, Table S4**).

**Table 3.**
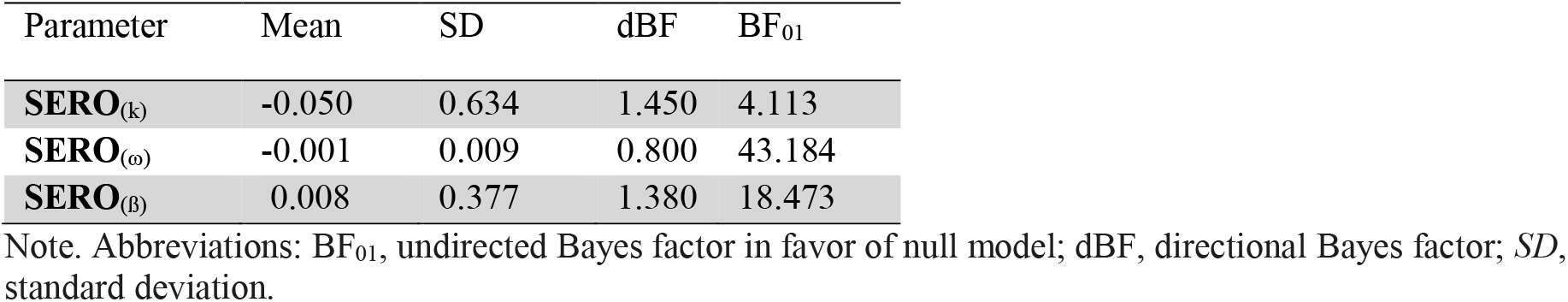
Summary statistics of the posterior distributions of computational shift-parameters (Offset-Model)

**Figure 8.**
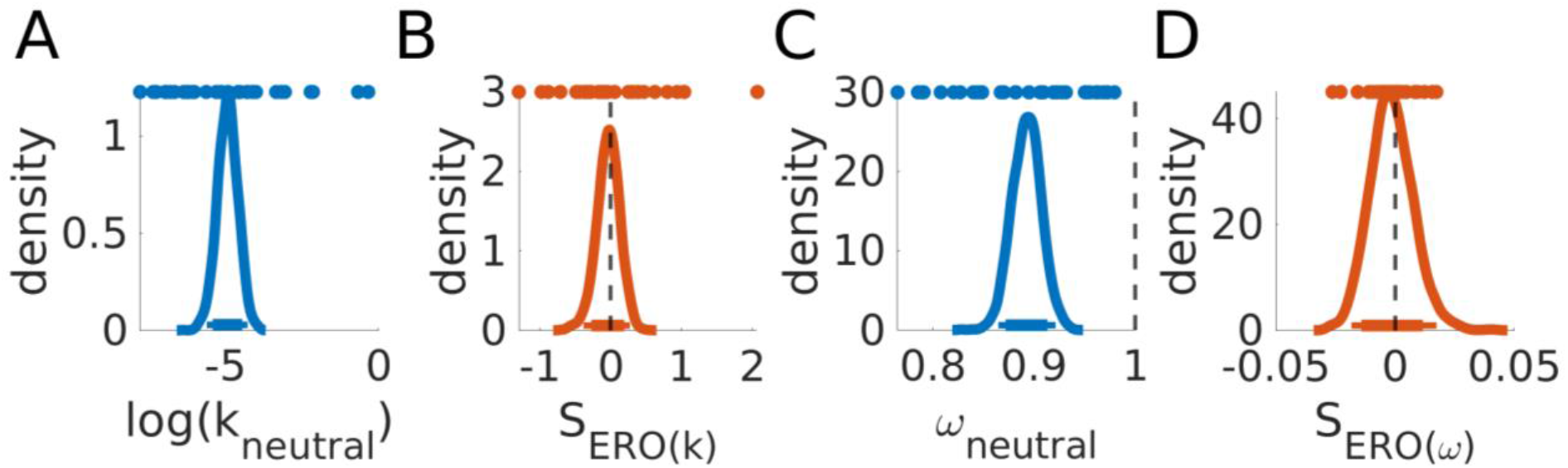
Posterior distributions for log(k_neutral_) and ω_neutral_ (A, C) as well as associated erotic shift parameters (SERO _(k, ω)_, B, D). Colored dots depict single subject posterior means. Thick and thin horizontal lines indicate 85% and 95% highest density intervals.

**Figure 9.**
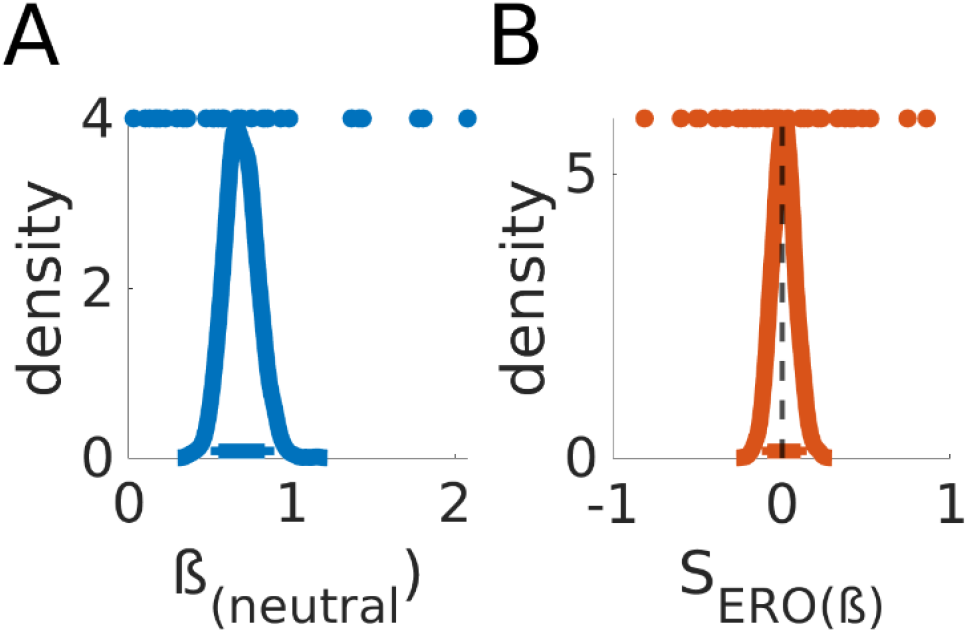
Posterior distributions for ß_neutral_ (A) and SERO_ß_ (B). Colored dots depict single subject means. Thick and thin horizontal lines indicate 85% and 95% highest density intervals.

To confirm the validity of our modeling approach, we also examined associations between *SEro*_(*k*)_ and model-free measures of TD (SS-option choice proportions). Correlations between model parameters and model-free measures were consistently in the expected direction (see **Figure S5**, **Supplementary materials**).

### 3.5. Appetitive cue effects on neuronal and behavioral indices of temporal discounting

Despite increased (sub-) cortical processing of erotic compared to neutral cues, TD did not differ between experimental conditions. We next assessed the preregistered links between neuronal cue-reactivity and TD. We first report cue exposure effects on DLPFC activity during LL-reward presentation (H5), possibly indicating changes in (prefrontal) cognitive control. We next show between-subjects associations between erotic reward-system-responsivity within key dopaminergic (Nacc, VTA) and prefrontal (DLPFC) areas, and changes in TD (H6).

Recall that we reasoned (and preregistered) that cue effects on TD reported in previous studies (Kim & Zauberman (2013); Van den Bergh et al. (2008); Wilson & Daly (2004)) might be due to cue-induced changes in prefrontal control regions and subcortical reward circuits. We tested the first prediction by comparing (left) DLPFC activity during LL-reward presentation (duration = 2s) between experimental conditions (H5) using the preregistered DLPFC mask and small volume correction (12mm sphere, peak coordinates (x = -38, y = 38, z = 6); Smith et al., 2018). Contrary to our hypothesis, we found no differences in DLPFC activity for the contrasts erotic > neutral or neutral > erotic. Likewise, on the whole brain level no voxels survived FWE (*p*<0.05) correction. A t-map depicting all activations associated with erotic > neutral LL-reward processing is publicly available at OSF (https://osf.io/9uzm8/).

Next, we tested associations between neuronal cue-reactivity-responses within key dopaminergic (Nacc, VTA) and prefrontal (DLPFC) areas and subject-specific condition effects on behavior (SERO_(k),_ SERO_(ω),_ see H6), capturing individual differences of cue effects (H6). Associations were quantified via Bayesian correlations (using JASP) separately for peak voxels from preregistered subcortical (Nacc, VTA) and cortical (DLPFC) ROIs (see methods section for details). We found no evidence for a correlation between functional cue-reactivity towards erotic cues and change in discounting behavior (SERO_(k),_ SERO_(ω);_ see **Figure 10)**.

**Figure 10.**
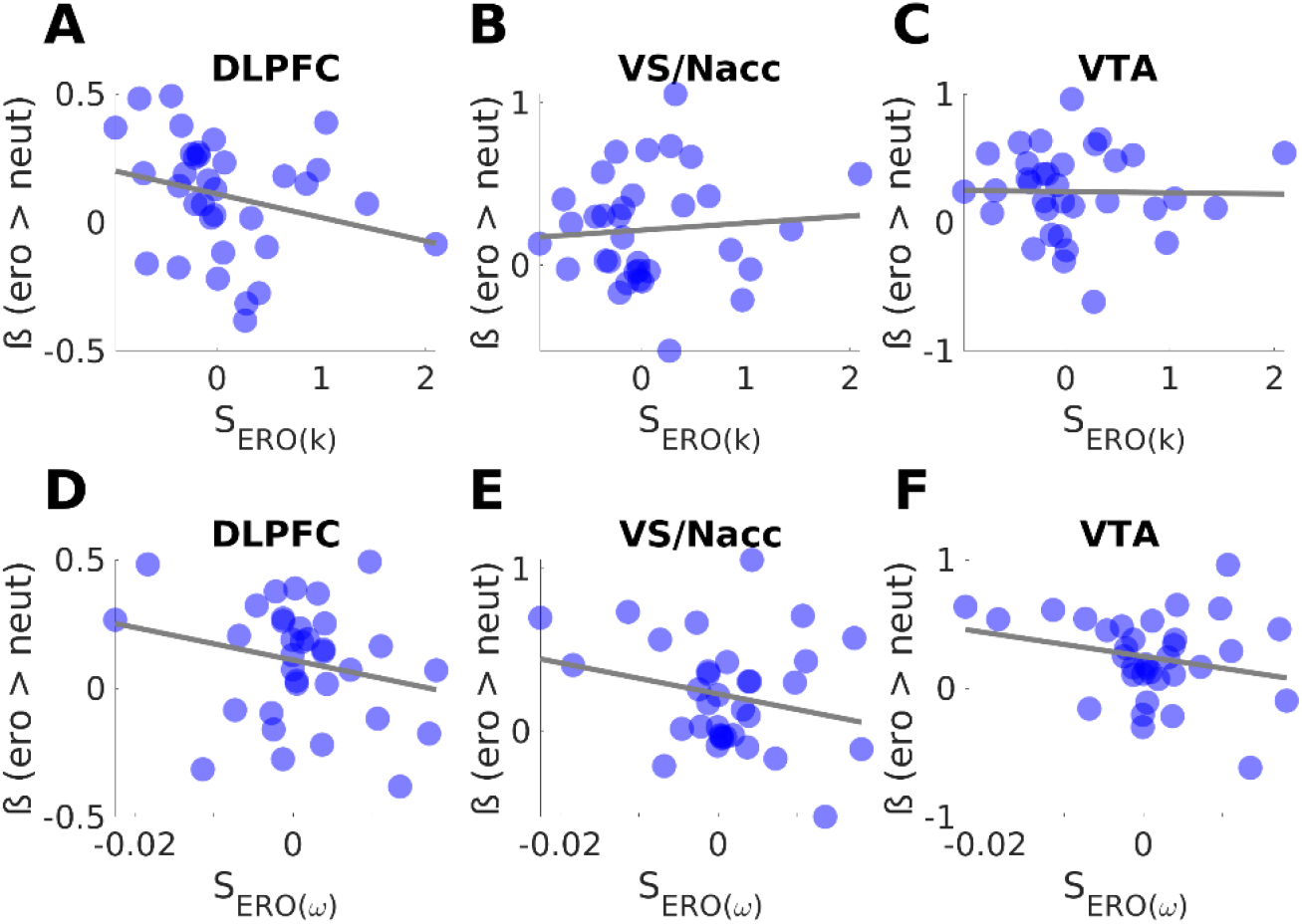
Associations between neuronal cue-reactivity-responses within key dopaminergic (VS/Nacc, VTA) and prefrontal (DLPFC) areas and subject-specific shift-parameters (S_Ero(k),_ S_Ero(ω)_)

Contrary, associations between cue evoked changes in TD (SERO_(k)_) and subcortical ROI activity (Nacc, VTA) yielded highest BF01 (Nacc: 4.226; VTA: 4.663). This indicates, we found at least moderate evidence for a model assuming no association between dopaminergic brain activity and changes in steepness of TD, and this model was approximately 4 to 4.5 times more likely than an alternative model given the data (see **Table 4**).

**Table 4.** Correlation statistics quantifying associations between peak voxel activity in key dopaminergic (VS/Nacc, VTA) and prefrontal (DLPFC) areas and subject-specific shift-parameters (S_ERO(k),_ S_ERO (ω)_) at the subject level.

| ROI<br>peak voxel [x,y,z] | Model parameter | Correlation<br>coefficient(r) | CI | BF01 |
| --- | --- | --- | --- | --- |
| **DLPFC [-30, 36, 4]** | $S_{ERO(k)}$ | -0.259 | [-0.531, 0.085] | 1.638 |
| | $S_{ERO(\omega)}$ | -0.239 | [-0.516, 0.105] | 1.932 |
| **VS/Nacc [-10, 2, -10]** | $S_{ERO(k)}$ | 0.083 | [-0.252, 0.393] | 4.226 |
| | $S_{ERO(\omega)}$ | -0.242 | [-0.518, 0.102] | 1.881 |
| **VTA [-10, 0, -12]** | $S_{ERO(k)}$ | -0.018 | [-0.340, 0.309] | 4.663 |
| | $S_{ERO(\omega)}$ | -0.240 | [-0.517, 0.104] | 1.911 |

Notes. ROI: Region of interest; CI: 95%-confidence interval; BF<sub>01</sub>, undirected Bayes factor in favor of null model

## 4. Discussion

Here we followed up on the literature on erotic cue exposure effects on TD (Kim & Zauberman, 2013; Mathar et al., 2022; Van den Bergh et al., 2008; Wilson & Daly, 2004). We expanded previous work by leveraging a preregistered fMRI approach to assess cue exposure-related activity changes in prefrontal and subcortical reward-related brain areas, and by linking these effects to TD. We first replicated a range of effects from the imaging literature on TD, including subjective value coding in vmPFC, striatum and cingulate cortex (Peters & Büchel, 2009) and increased left DLPFC activity for LL vs. SS choices (Smith et al., 2018). We also replicated the finding of increased visual and subcortical reward-related responses for erotic vs. neutral cues (Gola et al., 2016; Markert et al., 2021; Stark et al., 2019; Wehrum-Osinsky et al., 2014). However, these effects did not lead to increased TD, neither overall, nor in preregistered between-subject correlations focusing on key dopaminergic (Nacc, VTA) and prefrontal regions (DLPFC).

### 4.1. Neuronal correlates of subjective value and choice

We preregistered two replications for neural effects underlying TD. As predicted, and in line with previous work, activity in vmPFC, striatum, and cingulate cortex tracked subjective discounted value (SV) of LL-options (Bartra et al., 2013; Clithero and Rangel, 2013; Lee et al., 2021; Levy and Glimcher, 2012; Peters & Büchel, 2009; Sescousse et al., 2013). This effect was generally observed in most subjects and similarly evident following neutral and erotic cue exposure (at least for VMPFC and striatum). We found no evidence for condition differences in any of the reported clusters. This observation is inconsistent with the idea that upregulated activity levels e.g., in (dopaminergic) striatal regions following erotic cue exposure might disrupt subjective value encoding, which in turn might promote impulsive responding (Miedl et al., 2014).

We then focused on (left) dorsolateral prefrontal cortex (DLPFC), a region frequently implicated in TD (Guo & Feng, 2015; Hare et al., 2014) and self-control more generally (Hare et al., 2009). As preregistered, we observed increased decision-related left DLPFC activity for LL vs. SS choices. This pattern was observed across both experimental conditions (neutral, erotic), with no evidence for condition differences. Elevated DLPFC activity during LL choices (Smith et al., 2018) might be due to increased cognitive control during LL selections. This is supported by (1) increased TD following DLPFC disruption (Figner et al., 2010) and (2) fatigue effects manifested in increased TD that were associated with reduced DLPFC excitability (Blain et al., 2016). Our preregistered analyses therefore confirm an involvement of DLPFC specifically in LL choices.

On the whole brain level, two additional areas, right insular cortex and a cerebellar cluster showed increased activity for LL vs. SS choices. Whereas cerebellum has been observed in a wide range of tasks involving cognitive control and inhibition processes (Bellebaum & Daum, 2007; D’Mello et al., 2020; Stoodley & Schmahmann, 2009), insula activity was found to be specifically activated in LL-reward decisions and to depict a critical brain area involved in delaying gratification (Wittmann et al., 2007). This also resonates with findings from previous studies, reporting changes in insular activation in people with deficient foresight (Tsurumi et al., 2014), or reduced bilateral insula volumes in pathological gamblers compared with healthy controls (Mohammadi et al., 2016).

### 4.2. Appetitive cues affect neuronal reward circuitry

Exposure to appetitive visual cues, presented in a *blockwise* manner can increase impulsive choice in subsequent TD tasks (Kim & Zauberman, 2013; Van den Bergh et al., 2008; Wilson & Daly, 2004). We reasoned such cue effects on TD to be at least in part driven by upregulated reward circuitry (Li, 2008; Stark et al., 2019; Yeomans & Brace, 2015), an account not directly tested before. We focused on predefined ROIs previously associated with erotic stimulus processing (Stark et al., 2019) and presented participants with 40 intact experimental (neutral, erotic) and 20 scrambled control images. A comparison of intact vs. scrambled visual image processing confirmed highly plausible activation patterns, including large clusters across occipital cortices and the entire visual stream (Margalit et al., 2017).

Exposure to (intact) erotic compared to (intact) neutral stimuli revealed increased activity in widespread cortical and subcortical brain areas. Preregistered ROI analysis (FWE_SVC_ < 0.05) yielded strong posterior occipital and temporal clusters showing increased cortical responses to erotic vs. neutral cues. However, subcortical effects in reward-related circuits (e.g., ventral striatum, vmPFC) in our data in many cases fell just beyond the ROI mask constructed from the Stark et al. (2019) data, which mainly contained more dorsal striatal effects. We therefore followed up with an additional ROI analysis that used the same reward mask that we used (and preregistered) for the subjective value analysis (bilateral striatum, vmPFC, PCC and ACC) based on two meta-analyses (Bartra et al., 2013; Clithero & Rangel, 2014)). This confirmed significant bilateral activations in ventral striatum and VMPFC.

Our results are consistent with previously reported erotic cue responses across stimulus types (images or videos) and sexes (Ferretti et al., 2005; Mitricheva et al., 2019; Stark et al., 2019). While effects in parietal and occipital cortices might reflect attentional orientation towards erotic vs. neutral stimuli, striatal and anterior cingulate effects might reflect the intrinsic value of erotic vs. neutral cues (Georgiadis and Kringelbach, 2012; Kuehn & Gallinat, 2011; Poeppl et al., 2016a; Stark et al., 2019; Stoléru et al., 2012).

Neuronal cue-reactivity responses in visual regions largely overlapped with our preregistered ROI (based on group-level results (t-map) for the contrast erotic > neutral provided by Stark and colleagues (2019)). However, subcortical effects (e.g., in striatal regions) fell beyond the effects in the Stark et al. mask, and were instead located more ventrally, overlapping with the reward mask provided by the Rangel lab that we also used for the subjective value effects. We applied a binarization threshold (t-value = 6) to the entire T-map provided by Stark et al., to extract target voxels showing increased responsiveness to visual erotic stimuli. However Stark et al. (2019) used a somewhat longer stimulus duration (8s vs. 6s) and presented participants with both pictures and video clips to compare erotic vs. neutral cue reactivity responses. In their statistical analysis they did not differentiate between both stimulus types to increase generalizability. Stark et al. also used an expectation/anticipation phase prior to image/video onset which cued the nature of the upcoming stimulus (erotic or neutral). These differences might have contributed to the somewhat more ventral striatal effects that we observed compared to Stark et al. (2019).

### 4.3. No evidence for temporal discounting changes following blockwise exposure to appetitive cues

We used model-free and model-based approaches to quantify TD. Whereas model-free analyses focused on raw choice proportions, our best-fitting computational model allowed us to separate cue effects on steepness of TD (log(k)) from a delay-independent offset in the discounting curve. H1 was not confirmed - TD measures were not differentially affected by erotic cue exposure. Instead, Bayesian statistics suggested moderate evidence for the null model. This contrasts with earlier findings reporting increased TD following blockwise exposure to erotic visual stimuli (Kim & Zauberman, 2013; Van den Bergh et al., 2008; Wilson & Daly, 2004). On the other hand, it is consistent with a recent study from our group (Mathar et al., 2022) that used a similar cue exposure design. In Mathar et al. (2022), we used psychophysiology rather than fMRI. The lack of jitter between trial phases thus allowed us to use comprehensive modeling of RT distributions using diffusion models. Cue exposure led to a robust change in the starting point of the diffusion process towards SS options, but, as in the present study, did not reliably affect log(*k*).

Multiple reasons could account for this discrepancy. First, we used fMRI to assess neuronal correlates of cue-exposure and TD. The scanning environment, including loud noises, narrowness and movement restrictions itself might have acted as an external stressor, possibly attenuating behavioral effects. Indeed, neuroendocrine stress parameters (salivary alpha amylase, cortisol) increase at the beginning of an fMRI session (Gosset et al., 2018; Lueken et al., 2012; Muehlhahn et al., 2011), irrespective of stimulus presentation, and especially in scanner naïve participants (Tessner et al., 2006). Similarly, behavioral priming studies report smaller effects inside the scanner (Hommel et al., 2012), although such findings need replication. Both aspects might have contributed to an attenuation of behavioral cue effects in the current study. But, as noted above, in our earlier study (Mathar et al., 2022), cue exposure effects on log(k) where similarly largely absent, despite the lack of fMRI environment effects.

Further, our implementation of the cue-exposure phase differed slightly from previous approaches. Our cue phase was prolonged and included more experimental visual stimuli (n=40) than earlier studies (max n = 25; Kim & Zauberman, 2013; Mathar et al., 2022; Van den Bergh et al., 2008; Wilson & Daly, 2004), although this should arguably have increased behavioral effects. We included additional design changes due to the fMRI design (scrambled control images, attention checks, jitter intervals between stimuli). These aspects could have attenuated the *continuous* blockwise character of cue-exposure, and concomitant rise in tonic dopaminergic tone, which might be required to affect TD (Pine et al., 2010). This resonates with previous studies showing that intermittent/trialwise exposure to erotic cues is not sufficient to elevate TD (Knauth & Peters, 2022; Simmank et al., 2015).

Participants in our study passively viewed the presented images, rather than performing explicit arousal or valence ratings. However, explicit ratings might have induced deeper processing in earlier studies, which could have exhibited stronger effects on choice behavior (Van den Bergh et al., 2008; Wilson & Daly 2004). Such attention effects can modulate behavioral (Gawronski et al., 2010) and neural effects (Anderson et al., 2003) of emotional stimuli. However, passive versus active viewing of emotional images leads to similar physiological arousal effects (Snowden et al., 2016). Furthermore, our observation of increased activity in widespread cortical and subcortical networks in response to erotic vs. neutral control stimuli strongly argues against the idea that these cues were not adequately processed.

Taken together, behavioral effects of erotic cue exposure on TD might not be as unequivocal as previously thought (Kim & Zauberman, 2013; Van den Bergh et al., 2008; Wilson & Daly, 2004). Recent studies utilizing trialwise erotic cue exposure failed to find changes in TD (Simmank et al., 2015). More critically, cue-evoked physiological arousal did not predict changes in discounting behavior (Knauth & Peters, 2022), casting doubt on the idea of an upregulated internal arousal state, that drives approach behavior towards immediate reward (Knauth & Peters, 2022). Also, recent *blockwise* studies question simple main effects of erotic cue exposure on impulsivity. Some studies find that cue exposure effects only occur under specific motivational or metabolic conditions (e.g., hunger; Otterbring et al., 2020). A noted above, we recently observed a robust change in the starting point of the evidence accumulation process towards SS rewards, which was revealed by extensive drift diffusion modeling of response time distributions (Mathar et al., 2022), whereas log(k) was largely unchanged. It is thus possible that the detection of cue exposure effects might require modeling of choices *and* response times. However, our fMRI-based experimental design separated option presentation responses, thereby precluding us from using comprehensive diffusion modeling of response times.

### 4.4. Elevated activity levels in dopaminergic brain areas cannot account for behavioral changes in temporal discounting

A major strength of the current study is its ability to empirically test the theoretical assumption of a cue-evoked upregulation in neural reward circuits, which might reflect increased dopaminergic activity (O’Sullivan et al., 2011; Redouté et al., 2000). Such effects might facilitate reward approach across domains (Van den Bergh et al., 2008). This idea is supported by pharmacological modulations of central dopamine transmission that affect TD (Arrondo et al., 2015; Cools, 2008; de Wit, 2002; Hamidovic et al., 2008; Kayser et al., 2012; Petzold et al., 2019; Pine et al., 2010; Wagner et al., 2020; Weber et al., 2016).

Here, we directly examined associations between neuronal cue-reactivity-responses towards erotic cues within key dopaminergic (Nacc, VTA) and prefrontal (DLPFC) areas and subject-specific condition effects on TD (SERO_(k),_ SERO_(ω)_). However, our preregistered associations could not be confirmed.

While a general dopaminergic impact on TD is well established, direction of reported effects in human studies appears somewhat inconsistent. Pine et al. (2010) observed increased TD following administration of the catecholamine precursor L-DOPA vs. placebo in a small sample of n=14. In contrast, Petzold and colleagues (2019) observed no overall effect of L-DOPA administration on TD. Instead, effects depended on baseline impulsivity, supporting the view of an inverted-U-model of dopamine effects on cognitive control (Cools and D’Esposito, 2011). We recently observed (Wagner et al., 2020) reduced TD after a single low dose of the D2 receptor antagonist haloperidol, which is thought to increase striatal dopamine. The current study complements these previous findings and attempted to link (dopaminergic) reward system activity - which pharmacological approaches aim to evoke - to behavioral effects. However, upregulated reward system activity appears to be not sufficient to evoke *behavioral* cue effects (see previous section).

In the light of these contradictory findings, future studies should consider additional factors possibly involved in previously reported effects on TD. On the physiological level, arousal-related enhancement of noradrenaline (NE) release may be one possible mechanism (Ventura et al., 2008). Previous studies indeed find increased pupil dilation following highly arousing cues (Aston-Jones and Cohen, 2005; Finke et al., 2017; Kinner et al., 2017; Knauth and Peters., 2022; Murphy et al., 2011). NE agonists have been found to affect several forms of impulsivity (Robinson et al., 2008) and to directly increase the preference for LL rewards (Bizot et al., 2011). Further, Yohimbine, an α_2_-adrenergic receptor antagonist that increases NE release reduced discounting in humans (Herman et al., 2019; Schippers et al., 2016). It appears highly plausible that (appetitive) cue-exposure will always affect both, noradrenergic and dopaminergic neurotransmitter systems.

### 4.5. Limitations

Our study has a few limitations that need to be acknowledged. First, we only tested male participants. Men and women might differ in neuronal responses to affective stimulus material and emotional processing (Bradley et al., 2001; Lithari et al., 2010; Wrase et al., 2003), although a recent meta-analysis found at most negligible sex differences in neural correlates of sexual arousal (Mitricheva et al., 2019). However, to extent generalizability of results, future studies should include participants from both sexes.

Second, we did not include an image rating task, capturing arousal, valence or related dimensions. Therefore, we cannot directly quantify subjective arousal associated individual cues. However, fMRI revealed substantial differences in neural responses to erotic vs. neutral cues in plausible brain regions implicated in attention and reward. Further, a pilot study in an independent sample confirmed that the applied stimulus material clearly modulated subjective arousal. Still, future studies might complement fMRI and task-based measures with self-reported arousal.

Third, we did not include an additional aversive cue condition to control for unspecific arousal effects. Previously reported erotic cue effects on TD might be at least partly be attributable to increased arousal, although aversive cue effects on TD likewise appear mixed (Cai et al., 2019; Guan et al., 2015; Knauth & Peters, 2022). Nonetheless, it would be interesting to assess whether neuronal measures of aversive cue processing are predictive for choices.

Lastly, on each trial, participants only viewed the LL option, whereas the SS reward was fixed and never shown on the screen, as done in numerous earlier studies (Peters & Büchel, 2009; Kable & Glimcher 2007). However, additionally displaying the smaller sooner reward (separated by a further jitter interval) could be interesting for two reasons. First, although we showed that value computations for LL rewards was largely unaffected by cue condition, neuronal representations of an *immediate* reward might have been affected by cue condition. Second, elevated dopamine tone might foster approach behavior towards rewards that appear spatially near or available. Only presenting one of two possible choice options instead of both (Guan et al., 2015) might have biased or even compensated cue effects.

### 4.6. Conclusion

Previous studies indicated that highly appetitive stimuli might increase TD behavior (Otterbring et al., 2020; Kim & Zauberman, 2013; Wilson & Daly, 2004). Cue-reactivity in reward-related circuits was suspected as potential mechanism underlying these effects (van den Bergh et al., 2008). Here we leveraged combined fMRI during both cue exposure and decision-making to link activity in reward circuits to changes in TD. We first replicated core neural effects underlying TD (value coding in vmPFC, striatum and posterior cingulate, LL-choice-related activity in DLPFC) (Bartra et al., 2013; Clithero & Rangel, 2014; Kable & Glimcher, 2007; Peters & Büchel, 2009; Smith et al., 2018). Further, we confirmed increased (sub-) cortical processing during erotic vs. neutral cue exposure in core regions of the reward circuit. However, our preregistered hypothesis of increased TD following erotic cue exposure was not confirmed. This resonates with recent findings from our lab, where such effects were only observed for the bias parameter in the drift diffusion model, and not for choice behavior per se (Mathar et al., 2022). Importantly, and in contrast to our preregistered hypothesis, activity in key reward regions (Nacc, VTA) did not predict changes in behavior. Our results cast doubt on the hypothesis that upregulated activity in the reward system is sufficient to drive myopic approach behavior towards immediately available rewards.

## Data availability

Raw choice data, STAN modeling code of our (best fitting) computational model (used for subsequent analyses) as well as fMRI T-maps underlying all depicted figures are publicly available on the Open Science Framework (https://osf.io/9uzm8/).

## Supplementary Information

**Supplementary Table S1.**
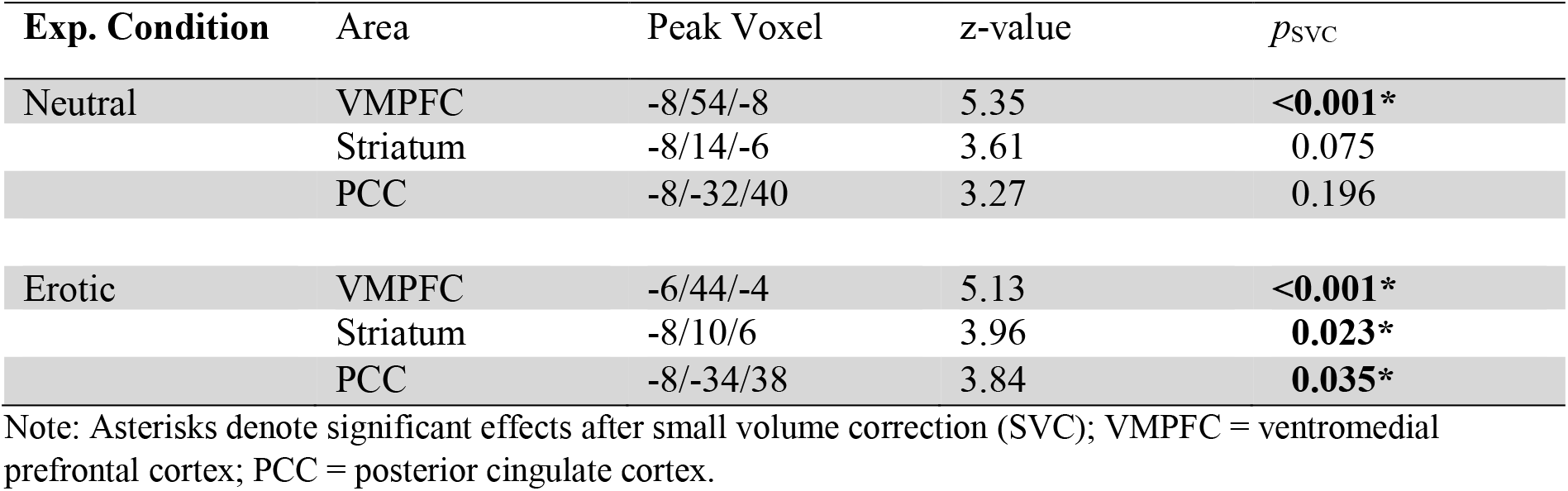
Peak voxels and SVC corrected *p*-values for condition-wise SV-coding (erotic, neutral)

**Supplementary Table S2.**
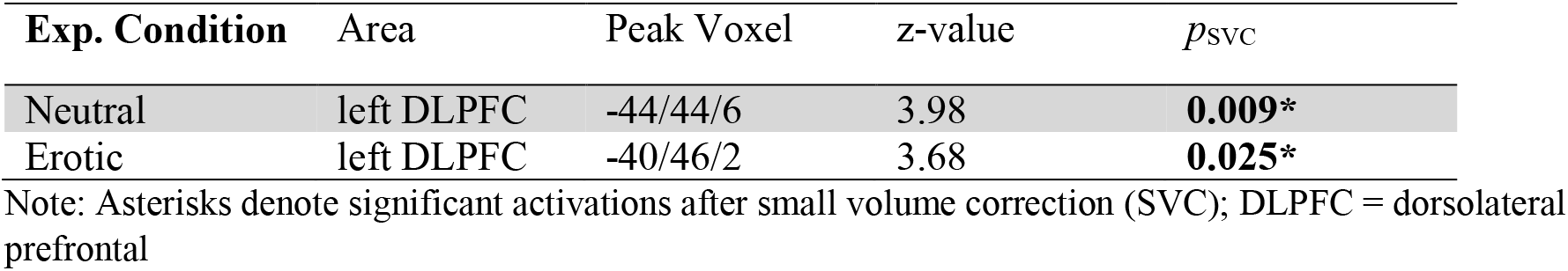
Peak voxels and SVC corrected *p*-values for condition-wise LL-choice coding (erotic, neutral)

**Supplementary Figure S1.**
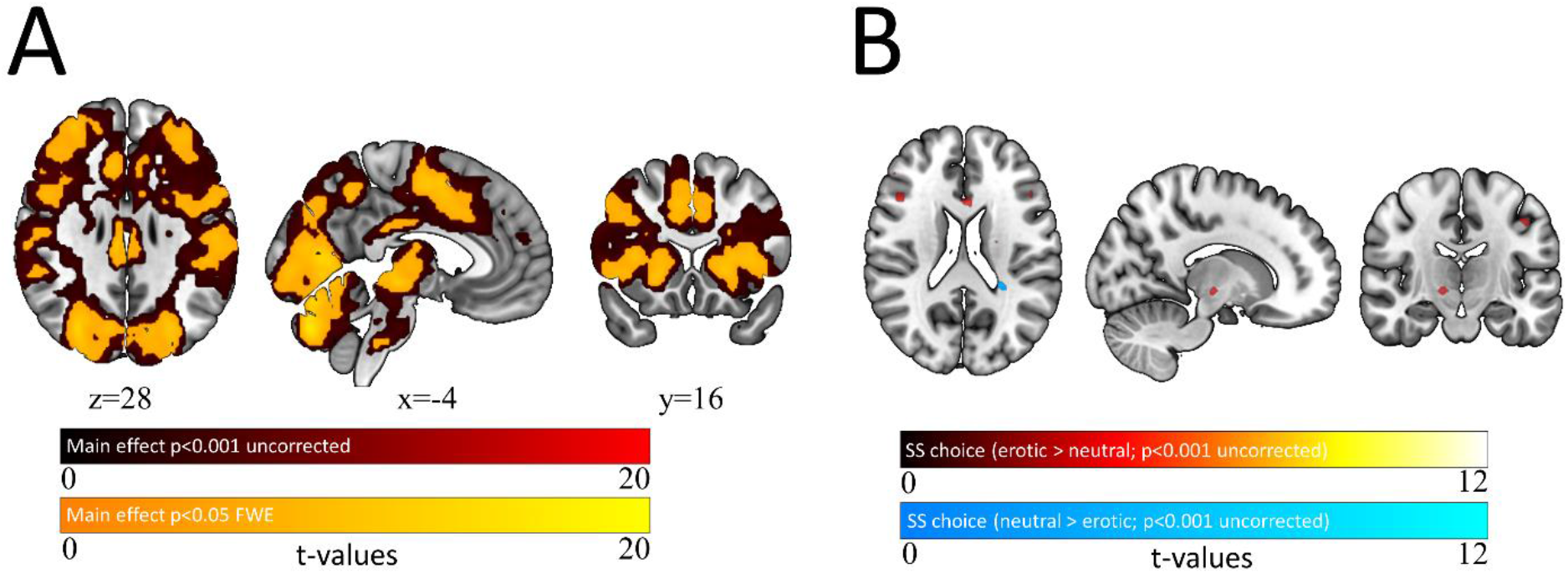
Neuronal correlates of smaller-sooner choices. **A**: Display of the parametric (SS-) choice-regressor (mean across conditions); red, *p*<0.001 (uncorrected); yellow, whole-brain FWE corrected *p* < 0.05; **B**: Condition contrasts, erotic > neutral (red/yellow) and neutral > erotic (light blue), p<0.001 uncorrected (whole brain analysis).

**Supplementary Figure S2.**
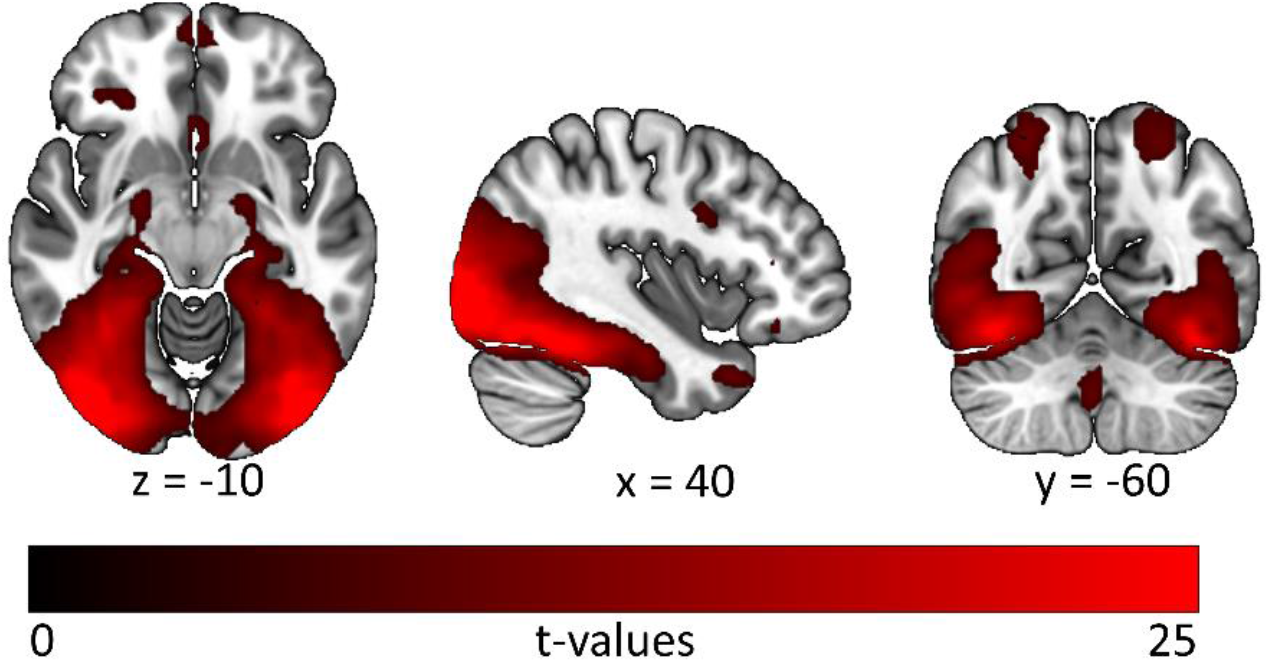
Neuronal correlates intact vs. scrambled image processing irrespective of experimental condition. Red, whole-brain FWE corrected *p* < 0.05 (whole brain analysis).

**Supplementary Table S3.** Proportions of correctly predicted binary choices (mean [CIs]) for both included TD models (Base-Model, Offset-Model) split by experimental condition (see also Supplementary Figure S3).

| Model | Neutral | Erotic |
| --- | --- | --- |
| Base-Model | 0.799 [0.762-0.836] | 0.793 [0.753-0.833] |
| Offset-Model | 0.831 [0.791-0.870] | 0.812 [0.766-0.858] |

**Supplementary Figure S3.**
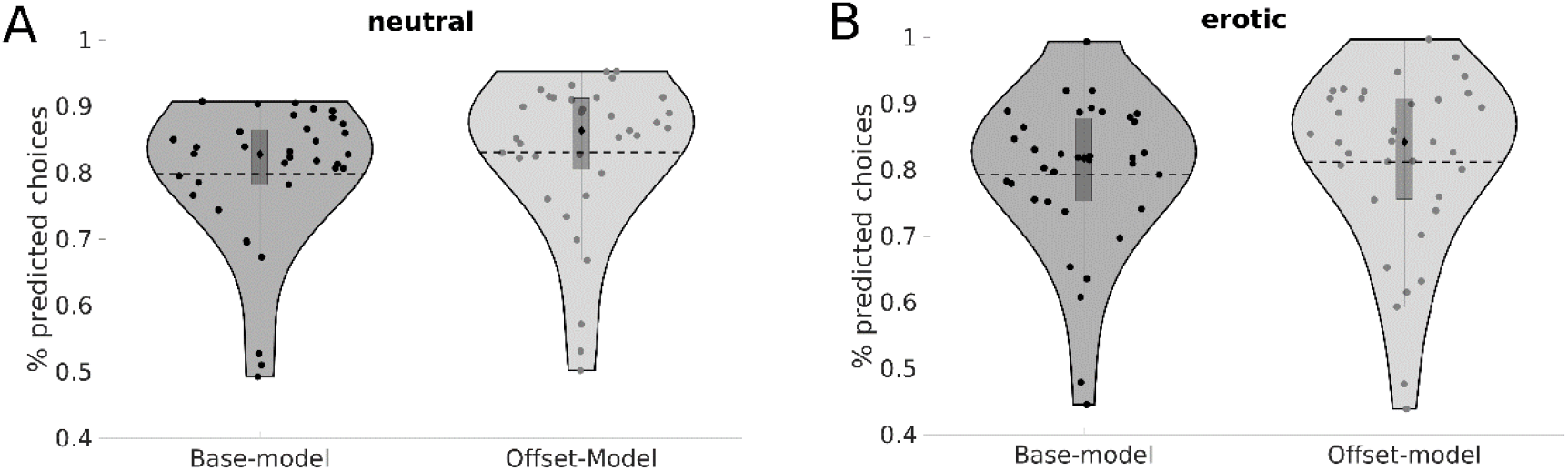
Proportions of correctly predicted binary choices for the Base-Model and the Offset-model (including an SV-offset parameter ω) split by experimental condition (A: neutral, B: erotic); Dots = single subjects; Dashed lines = group means; Black diamonds = group medians.

**Supplementary Figure S4.**
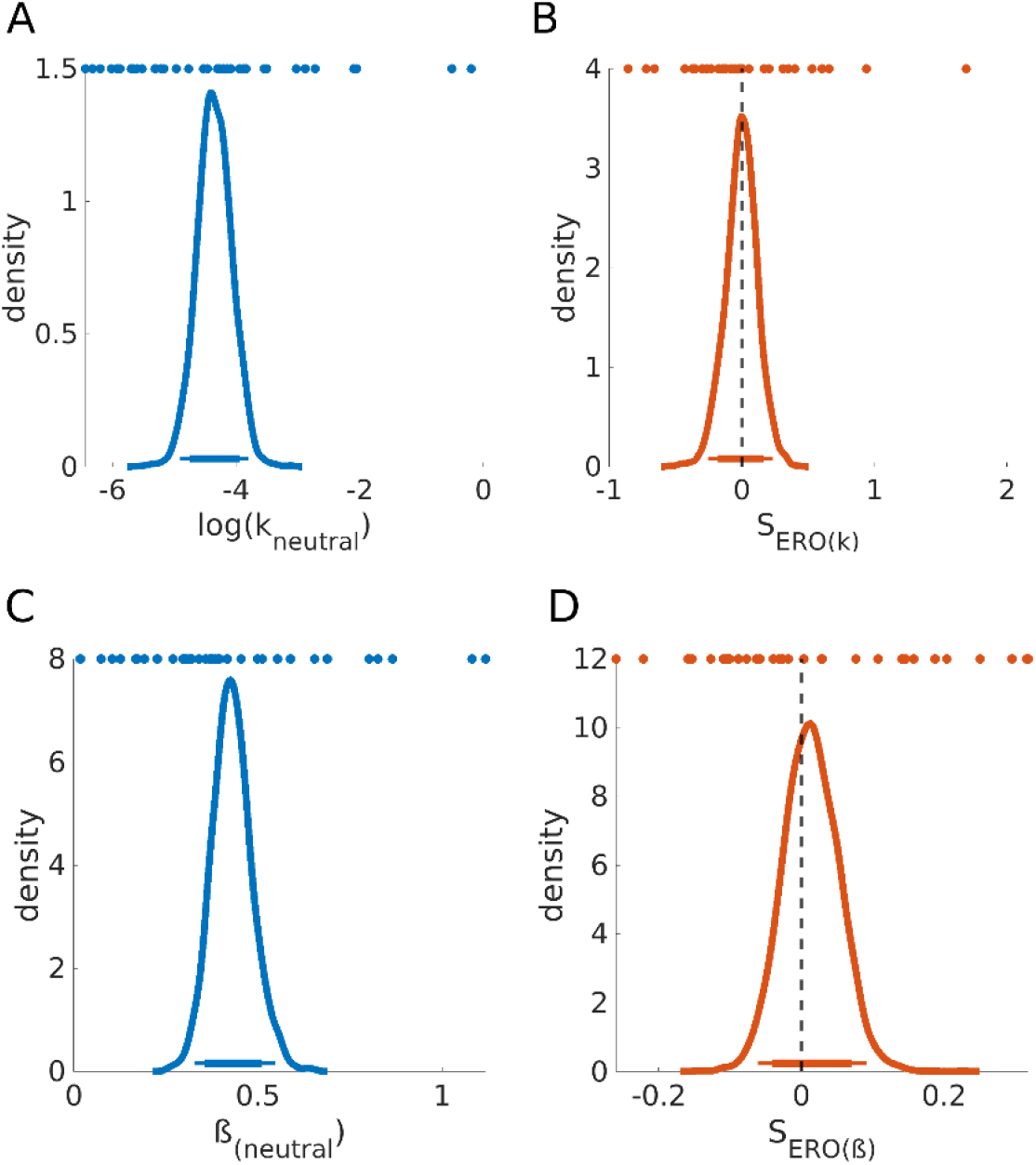
Posterior distributions for log(*k*_neutral_) (A), ß_(neutral)_ (C) and associated erotic shift parameters (B & D; Base-Model); Colored dots depict single subject means. Thick and thin horizontal lines indicate 85% and 95% highest density intervals.

**Supplementary Table S4.** Summary statistics of the posterior distributions of computational shift-parameters (Base-Model)

| Parameter | Mean | SD | dBF | BF<sub>01</sub> |
| --- | --- | --- | --- | --- |
| S<sub>Ero(k)</sub> | -0.004 | 0.484 | 1.030 | 5.810 |
| S<sub>Ero(β)</sub> | 0.014 | 0.154 | 0.530 | 30.807 |

Note. Abbreviations: BF<sub>01</sub>, undirected Bayes factor in favor of null model; dBF, directional Bayes factor; SD, standard deviation.

**Supplementary Figure S5.**
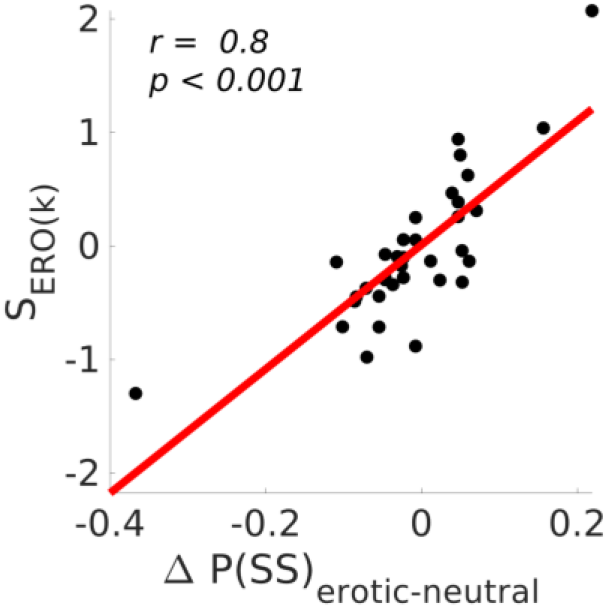
Associations between model-free (SS-choice proportions) and model-based measures (S_Ero(*k*)_) of temporal discounting behavior (Offset-model); *r* = Pearson’s correlation coefficient.

